# mTORC1-dependent SOCE activity regulates synaptic gene expression and muscle response to denervation

**DOI:** 10.1101/2024.04.01.587665

**Authors:** Alexandre Prola, Olivier Dupont, Jayasimman Rajendran, Florent Chabry, Stéphane Koenig, Maud Frieden, Perrine Castets

**Affiliations:** Department of Cell Physiology and Metabolism, Faculty of Medicine, University of Geneva, 1 rue Michel Servet, CH-1211 Geneva, Switzerland

**Keywords:** SOCE, NMJ, mTORC1, muscle denervation, synaptic gene, calcium micro-domains

## Abstract

Neuromuscular junction (NMJ) instability is central in muscle dysfunction occurring in neuromuscular disorders and aging. NMJ maintenance requires regionalized regulation of synaptic genes, previously associated with Ca^2+^-dependent pathways. However, what sustains Ca^2+^ micro-domains in myofibers and allows a rapid response to denervation is not known. Here, we identify that Store-Operated Calcium Entry (SOCE) plays a critical role in synaptic gene regulation. SOCE components show differential enrichment in sub- and non-synaptic muscle regions. Especially, STIM1 accumulation at rough endoplasmic reticulum associates with functional SOCE at the endplate. Denervation increases SOCE in non- and sub-synaptic regions, together with reticulum remodeling. *Stim1* knockdown hampers denervation-induced synaptic gene up-regulation, while STIM1 overexpression increases synaptic gene expression in innervated muscle. Finally, mTORC1 activation mimics the effect of denervation on SOCE capacity, STIM1 localization and reticulum remodeling. Together, our results reveal a decisive role of SOCE in sensing innervation and regulating muscle response to denervation. They further suggest that SOCE perturbation may contribute to neuromuscular integrity loss in pathological conditions associated with mTORC1 dysregulation.

## Introduction

Skeletal muscle contraction is governed by motor neuron at the level of the neuromuscular junction (NMJ), which defines a highly specialized sub-synaptic region in myofibers, called endplate. Synaptic proteins, including acetylcholine receptors (AChRs), are strictly expressed by sub-synaptic nuclei and aggregate at the endplate in innervated muscle ^1^. The establishment and maintenance of a unique synapse per myofiber rely on the electrical activity and subsequent calcium (Ca^2+^) transients in muscle ^2, 3^. Indeed, activity-dependent Ca^2+^ fluxes govern synaptic gene repression in non-synaptic nuclei ^4^, which would involve calcium/calmodulin-dependent protein kinase II (CaMKII)-dependent phosphorylation of myogenin and histone deacetylase 4 (HDAC4) in non-synaptic region ^5–7^. In parallel, at the endplate, Ca^2+^ fluxes drive agrin-induced AChR clustering ^8^ and AChR recycling ^9^, whilst preserving synaptic gene expression. This model suggests yet-unidentified mechanisms governing the establishment of specific functional Ca^2+^ micro-domains in non- and sub-synaptic regions of myofiber.

In response to nerve injury, myofibers release neurotrophic factors that favor sprouting of neighboring nerves and eventually muscle reinnervation ^10^. Poor reinnervation, caused by a defective muscle response to denervation, leads to muscle atrophy, a process involved in several neuromuscular diseases and systemic conditions, such as aging ^11, 12^. The local response of the sub-synaptic muscle region to denervation, which includes an increase in synaptic gene expression and AChR turnover, supports its reinnervation at the original endplate ^13–15^. In addition, non-synaptic myonuclei release synaptic gene repression. By extrapolation with activity-dependent processes, the response to inactivity is suggested to arise from the loss of Ca^2+^ transients and decreased CaMKII activity, resulting in myogenin- and HDAC4-dependent re-expression of synaptic genes ^16–18^. However, changes in Ca^2+^ fluxes and CaMKII activity have never been assessed in muscle after nerve injury.

Store-Operated Calcium entry (SOCE) is a tightly regulated Ca^2+^ influx that allows to refill intracellular Ca^2+^ stores. A reduction in Ca^2+^ levels in the endoplasmic reticulum (ER) triggers the oligomerization of the transmembrane proteins stromal interaction molecule 1 or 2 (STIM1/2), and their interaction with Ca^2+^ channels, such as Orai1, present at the plasma membrane. This induces cellular entry of Ca^2+^, which is then transported back to the ER by sarcoendoplasmic reticulum calcium ATPase (SERCA) pumps ^19^. The importance of SOCE in muscle is evident as mutations in *STIM1* or *ORAI1* cause muscle diseases ^20–22^. However, the physiological importance of this relatively small Ca^2+^ influx as compared to the large intracellular Ca^2+^ stores remains highly debated ^23–29^.

Here, we explored the role of SOCE in the regionalized expression of synaptic genes and the muscle response to denervation. We identify that STIM1 and Orai1 are enriched at the endplate, which is associated with high SOCE capacity. Muscle denervation increases SOCE, which may contribute to the release of synaptic gene expression. Consistently, STIM1 overexpression increases synaptic gene expression in innervated muscle, while STIM1 depletion hampers synaptic gene up-regulation upon denervation. Finally, constant activation of mammalian target of rapamycin complex 1 (mTORC1), a condition known to reduce muscle response to denervation ^30^, exacerbates SOCE. Together, our results reveal a yet-unknown role of SOCE in sensing neural activity in muscle and initiating the adaptive muscle response to denervation.

## Results

### Genes encoding SOCE components display regionalized expression in myofiber

To evaluate *in vivo* the role of Ca^2+^ fluxes and Ca^2+^-dependent signaling pathways in the muscle response to nerve injury, we cut unilaterally the sciatic nerve of adult mice to trigger denervation of hindlimb skeletal muscles. We then assessed CaMKII autophosphorylation on T286/287 residue as a marker of its kinase activity ^31^ in *tibialis anterior* (TA) muscle, 3 to 48 hours and 7 days after nerve injury, *i.e.,* before and after the peak of myogenin expression occurring 3 days after denervation ^32^. Autophosphorylation of CaMKII isoforms was unchanged 3 to 48 hours after denervation, and then increased in 7-days-denervated muscle compared to innervated muscle (Fig. 1A). In innervated TA muscle, the active form of CaMKII isoforms (CAMKII-P^T286/7^) accumulated at the NMJ, stained with α-bungarotoxin (BTX) that binds AChRs (Supplementary Fig. S1A). CAMKII-P^T286/7^ accumulation was unchanged in response to denervation (Supplementary Fig. S1A). These results suggest that denervation-induced synaptic gene expression does not rely on CaMKII inhibition. To identify alternative Ca^2+^-dependent pathways possibly involved in synaptic gene regulation, we screened for genes encoding Ca^2+^ regulators with regionalized expression, by comparing their transcriptional expression in non- *vs* sub-synaptic regions using SarcoAtlas database ^11^. Out of the 378 Ca^2+^-associated genes obtained from Gene Ontology database and detected in SarcoAtlas, 110 genes, including *Camk2d,* showed stronger expression in the sub-synaptic region (Supplementary Fig. S1B, C). Clustering of these genes based on the molecular function of the encoded proteins revealed that 76% of the genes associated with the regulation of or regulated by SOCE were preferentially expressed in the sub-synaptic region (Supplementary Fig. S1C, D). To confirm the regionalized expression of SOCE-associated genes, we next used single-nuclei RNA sequencing (snRNAseq) data on innervated TA muscle from adult mice publicly available in Myoatlas ^33^. Indeed, the genes encoding the main SOCE components STIM1, STIM2 and Orai1, were strongly expressed in sub-synaptic nuclei (Fig. 1B). Of note, expression of the genes encoding transient receptor potential canonical channel 3 (TRPC3) and Naked2, both recently associated with SOCE ^34, 35^, was predominant in sub-synaptic nuclei in myofibers (Fig. 1B and Supplementary Fig. S1E). Expression of *Orai2/3* and other *Trpc* genes was too low in these snRNAseq data to conclude on their expression pattern. Finally, *Camk2d* expression was higher at the myotendinous junction and in sub-synaptic nuclei, as identified in SarcoAtlas database (Supplementary Fig. S1E).

**Fig. 1.**
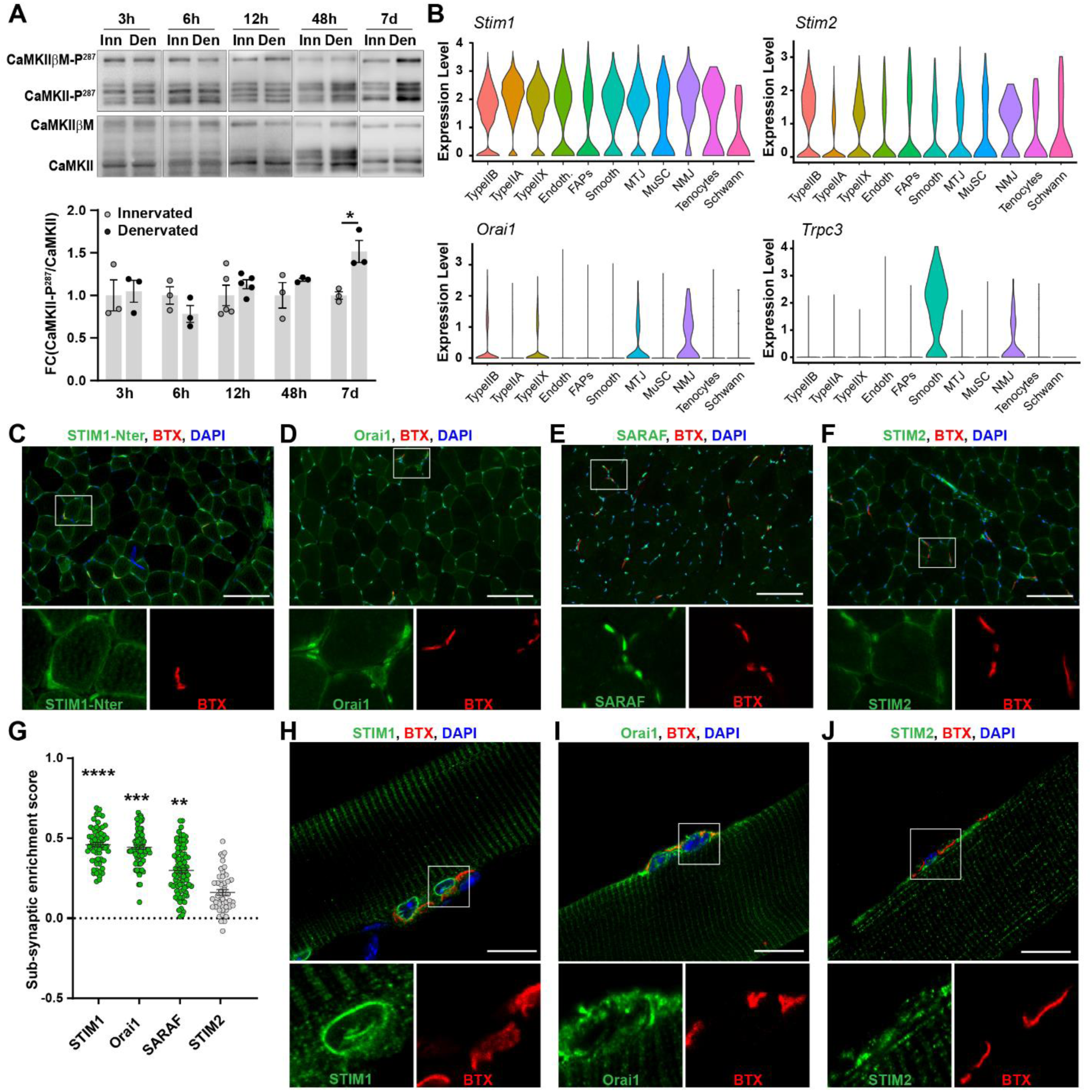
SOCE machinery displays regionalized localization in skeletal muscle fibers. (**A**) Representative immunoblots (n=3; except at 12hrs, n=5) of phosphorylated (Thr286/287) and total CaMKII in innervated (Inn) and denervated (Den) TA muscles, 3, 6, 12, 48 hours and 7 days after nerve injury. Levels of CaMKII-P^287^ are normalized to total CaMKII levels. (**B**) Violin plots showing expression levels of *Stim1*, *Stim2*, *Orai1* and *Trpc3* in clusters of single nuclei isolated from TA muscle of 3-month-old mice ^33^. Endoth: endothelial cells; FAP: Fibro/adipogenic progenitors; Smooth: smooth muscle; MTJ: myotendinous junction; MuSC: muscle stem cell. (**C-G**) Immunostaining of TA muscle sections for STIM1-Nter (**C**), Orai1 (**D**), SARAF (**E**) and STIM2 (**F**). The sub-synaptic region is visualized with α-Bungarotoxin (BTX). Scale bars, 100 µm. Sub-synaptic protein enrichment score is based on the quantification of the colocalization of proteins and BTX (**G**). Significantly enriched proteins are represented in green. Each dot represents a neuromuscular junction (> 40 NMJ from 4 independent samples). (**H-J**) Confocal images of STIM1-Cter (**H**), Orai1 (**I**), STIM2 (**J**) immunostaining with BTX staining of isolated EDL fibers. Scale bars, 20 µm. All data are mean± s.e.m.; *P < 0.05, **P < 0.01, ***P < 0.001, ****P < 0.0001; unpaired two-tailed Student’s t-test (A), P value based on t-score (G, see methods for more details).

To confirm the enrichment in SOCE machinery in myofiber sub-synaptic region, we immunostained TA sections for SOCE components, together with BTX (Fig. 1C-F), and calculated enrichment based on colocalization scores (Fig. 1G). Using 3 different antibodies targeting N- or C-terminal region of STIM1, we observed a dotty staining on transversal muscle sections that fits with the previously reported STIM1 localization at the sarcoplasmic reticulum (SR) ^29, 36^, as well as a staining surrounding myonuclei (Fig. 1C, and Supplementary Fig. S1F, G). Interestingly, there was a clear enrichment of STIM1 staining at the NMJ (Fig. 1G). On transversal muscle sections, Orai1 also showed an enrichment at the NMJ, as well as a dotty staining that was consistent with its localization at T-tubules ^29^ (Fig. 1D, G). Similarly, SARAF, which regulates STIM1 activity ^37^, accumulated preferentially around myonuclei and at the NMJ (Fig. 1E, G). In contrast, STIM2 immunostaining gave a dotty staining, but it accumulated neither at the periphery of myonuclei nor at the NMJ (Fig. 1F, G). To examine SOCE component localization with higher resolution, we performed immunostaining on fixed isolated fibers from *extensor digitorum longus* (EDL) stained with BTX. In myofibers, STIM1 showed typical striated staining consistent with its accumulation in terminal cisternae and longitudinal SR ^29, 36, 38, 39^. Moreover, as suggested by the immunostaining on muscle sections, it accumulated at the endplate of myofiber with a strong perinuclear staining (Fig. 1H). Similarly, Orai1 showed striated staining reminiscent of its enrichment at T-tubules, as well as an accumulation at the endplate of myofibers (Fig. 1I). Finally, STIM2 also showed striated patterns, but was not enriched at the endplate (Fig. 1J). Taken together, these results indicate that SOCE components are differentially expressed in sub- and non-synaptic muscle regions, which may sustain regionalized SOCE activity in myofibers.

### Denervation induces STIM1 re-localization at the endplate

As the regionalized SOCE distribution is compatible with a role in sensing muscle innervation/denervation, we examined if nerve injury induces changes in SOCE in muscle fiber. First, we quantified the expression of SOCE-related proteins in TA muscle in innervated or denervated conditions (12 hours or 7 days). Protein levels of the NMJ-enriched SOCE effectors, STIM1, Orai1 and SARAF, were unchanged after denervation (Fig. 2A-C). In contrast, protein levels of STIM2, which did not show enrichment at the NMJ, were unchanged 12 hours after denervation (Fig. 2A, B), but increased after 7 days (Fig. 2A, C). Of note, protein levels of TRPC3 and Naked2 also increased upon denervation (Fig. 2A, C). We next evaluated changes in the localization of these components after denervation, using immunostaining of muscle sections or isolated fibers stained with BTX. There was no major change in Orai1, SARAF and STIM2 localization detected on muscle sections 7 days after denervation (Supplementary Fig. S2A, B). In contrast, STIM1 staining increased in the sub-synaptic region of 7-days-denervated muscle (Fig. 2D and Supplementary Fig. S2B). On isolated fibers, the perinuclear staining of STIM1 observed at the endplate in innervated muscle was lost 7 days after denervation, resulting in a more diffuse STIM1 localization in the sub-synaptic muscle compartment (Fig. 2E). The proportion of fibers with a sub-synaptic perinuclear staining of STIM1 was strongly reduced 7 days after denervation (Fig. 2F). Together, these results point to an activity-dependent regulation of SOCE in muscle, with denervation-induced changes in the expression or localization of non- and sub-synaptic SOCE components.

**Fig. 2.**
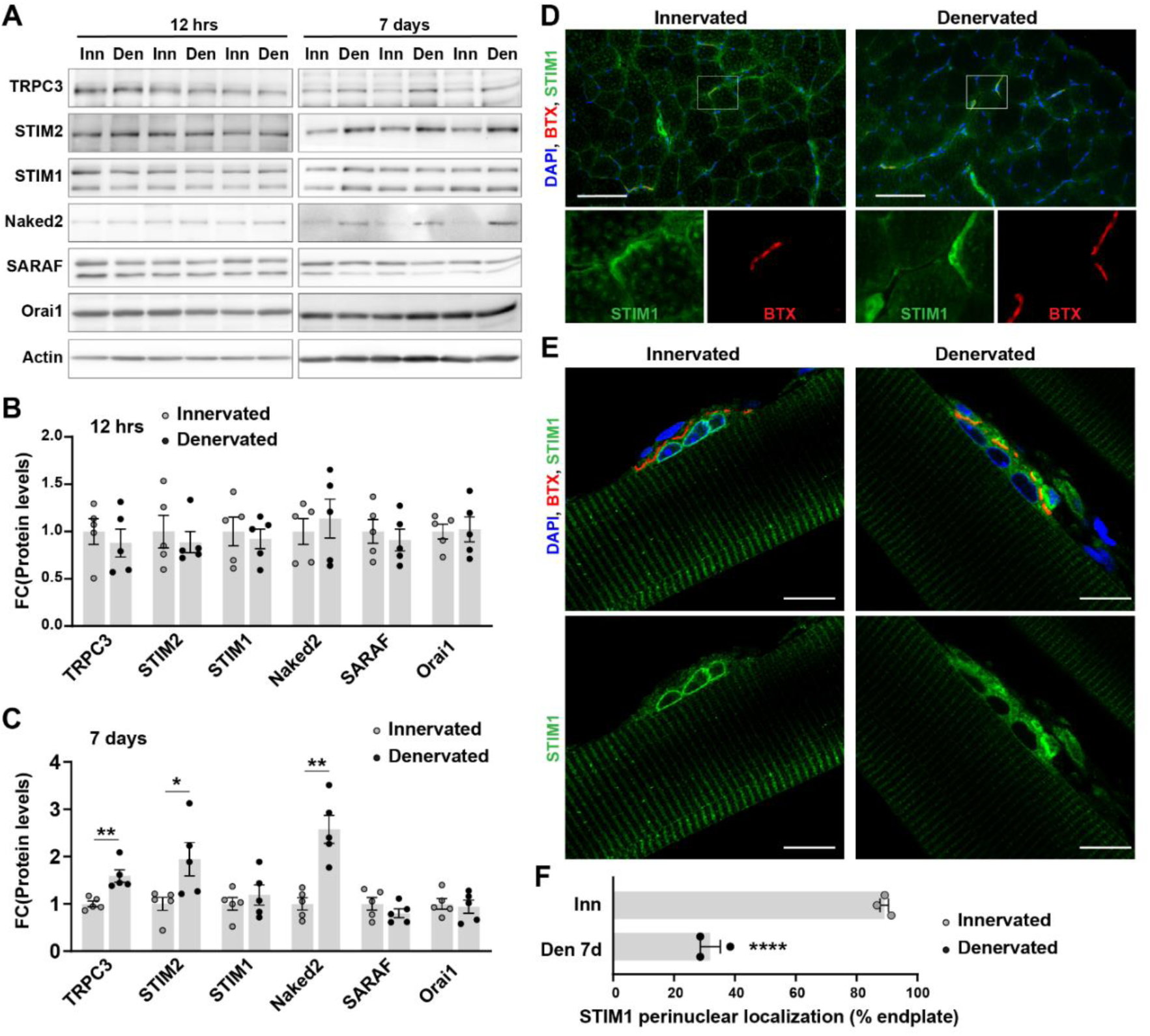
Denervation triggers STIM1 re-localization at the endplate. (**A-C**) Representative immunoblots of proteins involved in SOCE in innervated (Inn) and denervated (Den; 12hrs or 7 days) TA muscles. Quantification of protein levels, relative to Actin levels, is given in **B** (12hrs) and **C** (7 days). n=5. (**D**) Immunostaining for STIM1-Cter of innervated and denervated (7 days) TA muscle sections. The sub-synaptic region is visualized with α-Bungarotoxin (BTX). Scale bars, 100 µm. (**E, F**) Confocal images of STIM1-Cter immunostaining of fibers isolated from innervated or denervated (7 days) EDL muscles. Scale bars, 20 µm. The proportion of fibers with STIM1 perinuclear localization in the sub-synaptic region is given in **F**. n=3. All data are mean ± s.e.m; *P < 0.05, **P < 0.01, ****P < 0.0001; unpaired two-tailed Student’s t-test (B, C, F).

### rER enrichment and remodeling associate with activity-dependent SOCE

We hypothesized that STIM1 enrichment around myonuclei and at the endplate in innervated muscle corresponds to its accumulation at the rough endoplasmic reticulum (rER). Consistently, immunostaining of innervated TA sections against the rER marker, KDEL, showed major enrichment at the NMJ (Fig. 3A and Supplementary Fig. S3A), as opposed to the dotty staining of the SR marker, SERCA1 (Fig. 3B). In isolated fibers, we showed that STIM1 striated staining partially overlaps with SERCA1 staining, which was consistent with its previously reported localization in the longitudinal SR within the I-band region and in SR terminal cisternae ^29, 38–40^ (Fig. 3C). In parallel, STIM1 perinuclear staining, especially at the endplate, colocalized with KDEL staining (Fig. 3D). These results suggest that STIM1 accumulation in the sub-synaptic region of myofibers reflects an enrichment in rER in this region. To confirm the regionalized organization of reticulum network in muscle fibers, we used transmitted electron microscopy (TEM) to distinguish SR and rER in non- and sub-synaptic regions of EDL muscle. In non-synaptic region, SR was observed in the proximity of myofibrils, with longitudinal SR along sarcomeres and terminal cisternae juxtaposed to T-tubules, forming triads (Fig. 3E). In contrast, rER was rarely found in this region. Sub-synaptic regions were identified by the characteristic presence of folds of the sarcolemma, a cluster of sub-synaptic nuclei and mitochondria ^41^. There, we observed a large quantity of rER, very close or in direct contact with sarcolemma folds, as well as a high density of free ribosomes, consistent with high translational activity in this region ^30, 42^ (Fig. 3E). These results reveal a regionalized organization of the reticulum network in myofibers, which associates with STIM1 accumulation in SR and rER in non- and sub-synaptic regions, respectively.

**Fig. 3.**
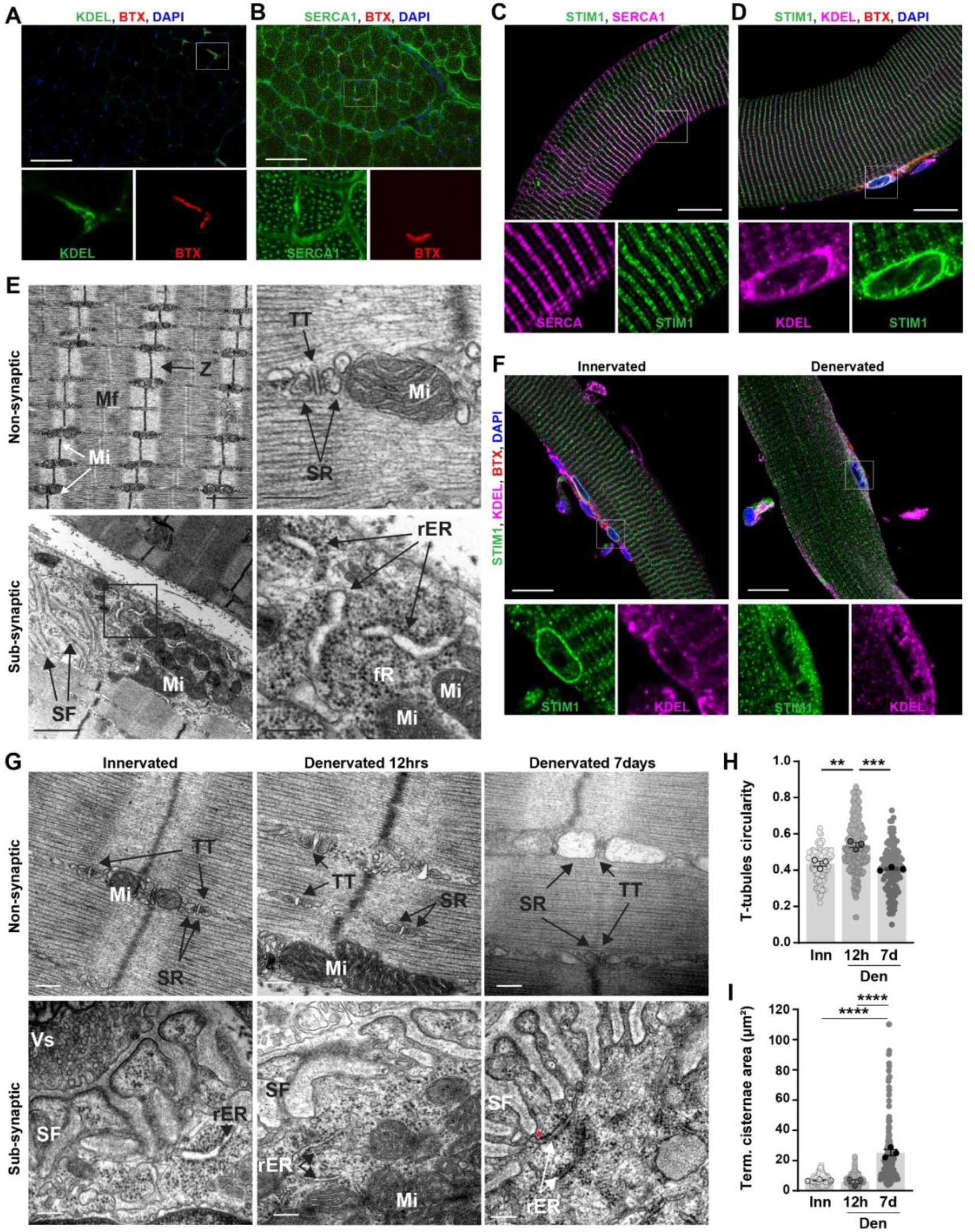
Denervation triggers remodeling of triads and rER. (**A, B**) Representative immunostaining of TA muscle sections for KDEL (**A**) and SERCA1 (**B**), with α-Bungarotoxin (BTX) staining. Scale bar, 100µm. Sub-synaptic protein enrichment score is provided in Supplementary Material, fig. S3A. (**C, D**) Confocal images of immunostaining for STIM1-Cter and SERCA1 (**C**) or KDEL (**D**) of isolated EDL muscle fibers. α-Bungarotoxin (BTX) is used to visualize the sub-synaptic region. Scale bar, 20 µm. (**E**) Representative images by transmission electron microscopy of longitudinal sections of myofibers from EDL muscle in the non- and sub-synaptic regions. The sub-synaptic region shows typical sarcolemma folds (SF) and a cluster of mitochondria (Mi). Higher magnification highlights the accumulation of rough endoplasmic reticulum (rER) and free ribosomes (fR). Z, Mf, TT and SR indicate a Z-line, myofibrils, a T-tubule and terminal cisternae of sarcoplasmic reticulum, respectively. Scale bars, 1 µm for low magnifications (left) and 100 nm for high magnifications (right). (**F**) Representative immunostaining for STIM1 and KDEL of isolated innervated and denervated (7 days) EDL fibers. BTX is used to visualize the sub-synaptic region. Scale bar, 20 µm. (**G**) Representative images of transmission electron microscopy of longitudinal sections of innervated and denervated (12 hrs or 7 days) EDL muscles in non- and sub-synaptic regions. Scale bars, 200 nm. TT, Mi, SR, Vs, SF, rER indicate T-tubules, mitochondria, sarcoplasmic reticulum, pre-synaptic vesicles, sarcoplasmic folds, rough endoplasmic reticulum, respectively. The red asterisk points to rER/sarcolemma contact points. (**H**) T-tubules circularity in innervated (Inn) and denervated (Den, 12 hrs or 7 days) EDL muscles based on electron micrographs. Light and opaque dots represent a T-tubule and the mean for one mouse, respectively. n=3. (**I**) Surface of SR terminal cisternae in innervated (Inn) and denervated (Den, 12 hrs or 7 days) EDL muscles based on electron micrographs. Light and opaque dots represent a terminal cisterna of SR and the mean for one mouse, respectively. n=3. All values are mean ± s.e.m; **P < 0.01, ***P < 0.001, ****P < 0.0001; one-way ANOVA with Benjamini and Hochberg’s post-hoc analysis to correct for multiple comparisons (H, I).

We next tested whether denervation-induced STIM1 re-localization reflects remodeling of rER in the sub-synaptic muscle region after nerve injury. Expression of SERCA1 and of the rER marker ribophorin 1 (RPN1) was similar between innervated and denervated muscles (Supplementary Fig. S3B). STIM1 colocalization with the rER marker KDEL, observed in innervated TA muscle, was preserved after 7 days of denervation (Supplementary Fig. S3C). Consistently, as observed for STIM1, the perinuclear staining of KDEL observed at the endplate in innervated fibers turned more diffuse after 7 days of denervation (Fig. 3F). This suggests that rER remodeling drives STIM1 relocalization in response to denervation. To further evaluate ultrastructural changes occurring in myofiber after denervation, we performed TEM on innervated or denervated (12 hours and 7 days) EDL muscles. The overall muscle ultrastructure was preserved 12 hours after denervation, with a regular arrangement of myofibrils and mitochondria (Supplementary Fig. S3D). Interestingly, the morphology of the triads was changed 12 hours after denervation, with a major alteration of the shape of T-tubules (Fig. 3G). Increased circularity (Fig. 3H) and roundness (Supplementary Fig. S3E) were consistent with a significant dilatation of T-tubules shortly after denervation. At this time point of denervation, there was no visible ultrastructural change in the sub-synaptic region (Fig. 3G and Supplementary Fig. S3D). After 7 days of denervation, the structure of the muscle started to be disrupted with the presence of misaligned sarcomeres (Supplementary Fig. S3D). Moreover, major dilatation of SR terminal cisternae, as reflected by the strong increase in their surface (Fig. 3I), led to the accumulation of large vesicles in 7-days-denervated muscle (Fig. 3G). Of note, the dilatation of T-tubules detected at 12 hours of denervation was transitory, as it normalized after 7 days of denervation (Fig. 3H and Supplementary Fig. S3D). In the sub-synaptic region, we observed an increase in the cytosolic space beneath sarcolemmal folds, with an accumulation of free ribosomes, rER, as well as lipid droplets 7 days after denervation (Fig. 3G and Supplementary Fig. S3D). Notably, close contacts between rER and sarcolemma that may sustain active SOCE were frequently observed at the endplate at this stage (Fig. 3G and Supplementary Fig. S3D). Together, these results indicate that denervation triggers sarcotubular remodeling in non-synaptic region, as well as rER remodeling in the sub-synaptic region that provides an ultrastructural basis for denervation-induced STIM1 relocalization at the endplate.

### SOCE capacity in sub- and non-synaptic muscle regions increases after denervation

To determine if STIM1/Orai1 enrichment in the sub-synaptic part of the muscle fiber translates into functional SOCE at the endplate, we first isolated live fibers from *flexor digitorum brevis* (FDB), loaded them with the Ca^2+^ dye Fura2-AM and stained them with BTX to discriminate non- and sub-synaptic regions (Fig. 4A). Freshly isolated fibers placed in Ca^2+^-free medium were then exposed to thapsigargin (TG) to inhibit SERCA pump and caffeine to trigger reticular Ca^2+^ efflux, followed by the addition of Ca^2+^ in the extra-cellular medium to potentially induce SOCE (Fig. 4B). In skeletal muscle fibers, full depletion of Ca^2+^ stores is unlikely to occur in physiological conditions. Similarly, experimental procedures used with myoblasts and myotubes to effectively deplete Ca^2+^ stores *in vitro* (such as TG and/or repeated caffeine treatments) are insufficient to fully deplete Ca^2+^ stores and/or rapidly provoke fiber contraction, even in the presence of the myosin inhibitor, N-benzyl-p-toluenesulfonamide (BTS) ^43–46^. However, a previous study reveals that local Ca^2+^ drop restricted to rER/SR micro-domains is sufficient to induce SOCE, even when Ca^2+^ stores are not fully depleted ^47^. Using similar procedure to induce partial Ca^2+^ store depletion in FDB fibers, we evaluated Ca^2+^ fluxes in two distinct regions of interest, defined based on BTX staining as to sub- and non- (> 30 µm from the endplate) synaptic regions. There was difference neither in cytosolic basal Ca^2+^ (Fig. 4C) nor in caffeine-induced Ca^2+^ release (Fig. 4D) between non- and sub-synaptic regions. Importantly, SOCE was detected in non- and sub-synaptic regions after adding Ca^2+^ in the medium, with similar SOCE amplitude in both regions (Fig. 4E, F). Of note, SOCE-dependent Ca^2+^ accumulation (measured as the area under the curve to the peak) was higher in the sub-synaptic region, as compared to non-synaptic region in these conditions (Supplementary Fig. S4A). These results demonstrate that the regionalized enrichment of SOCE components leads to functional SOCE throughout myofibers, including at the endplate.

**Fig. 4.**
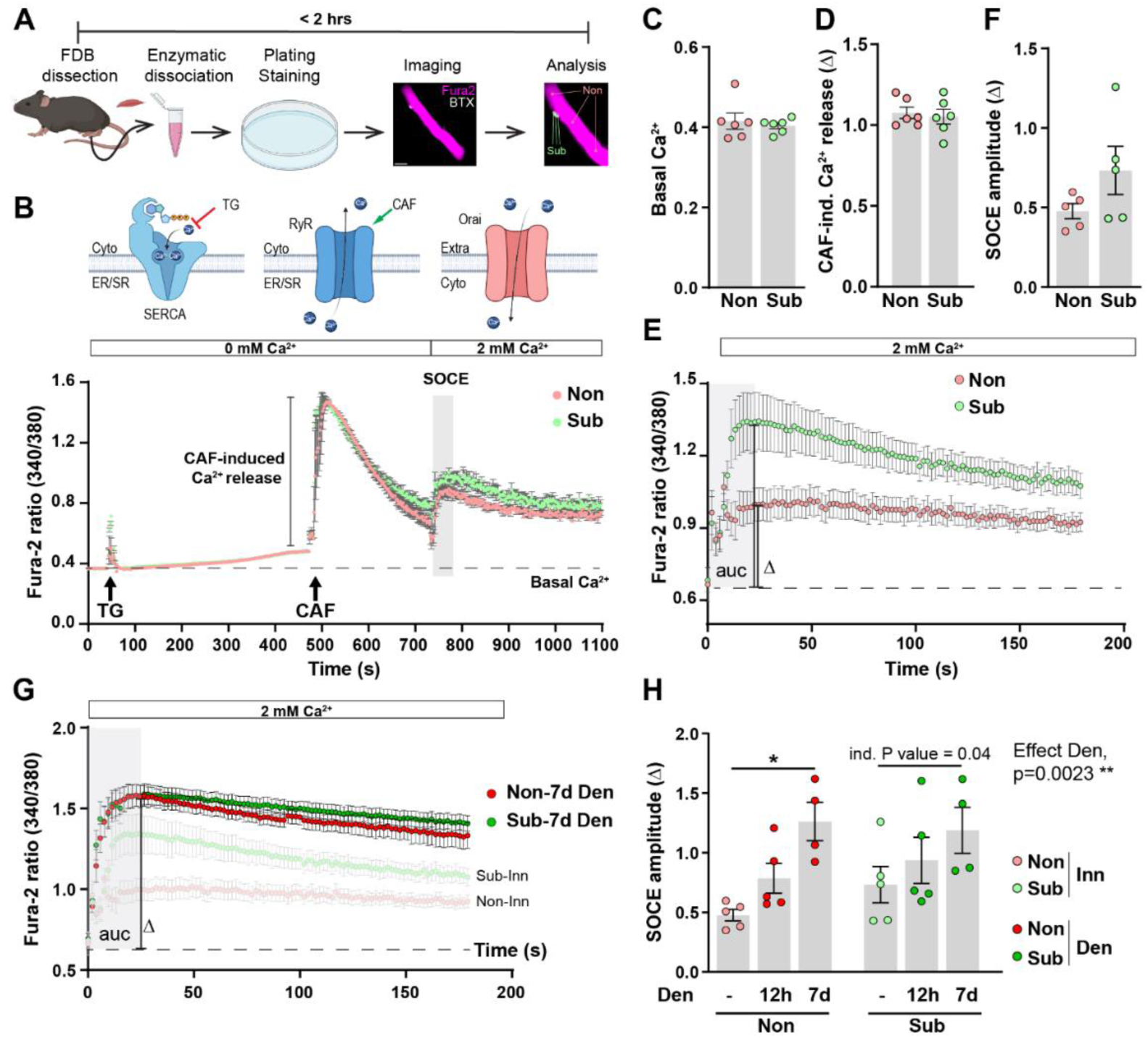
Regionalized SOCE capacity in non- and sub-synaptic muscle regions. (**A**) Workflow for FDB muscle fibers isolation, staining with Fura2-AM (purple) and BTX (white), and imaging. Scale bar, 50 µm. **(B)** Ca^2+^ fluxes measured in non- and sub-synaptic regions of FDB fibers after adding Thapsigargin (TG), Caffeine (CAF) or 2mM Ca^2+^ in the medium. (**C-F**) Cytosolic basal Ca^2+^ concentration (**C**), CAF-induced Ca^2+^ release (**D**) and SOCE amplitude (**F**) quantified in non- and sub-synaptic regions of muscle fibers. Mean SOCE traces in non- and sub-synaptic regions is shown in **E**. n=6 (C and D), 5 (F). (**G, H**) Mean SOCE traces (**G,** Den, 7 days) and SOCE amplitude (**H,** Den, 12hrs and 7 days) in non- and sub-synaptic regions of denervated FDB muscle fibers. In **G**, light traces correspond to innervated fibers (Inn). n=5 (Inn and Den, 12hrs), 4 (Den, 7 days). All data are mean ± s.e.m; *P < 0.05, **P < 0.01; two-way ANOVA with Benjamini and Hochberg’s post-hoc analysis to correct for multiple comparisons (H). Ind.: individual P value.

As previous reports propose that rER Ca^2+^ stores are easier to deplete than SR Ca^2+^ stores ^43–45^, SOCE detected at the endplate may arise in part from the enrichment in rER and SOCE machinery in this region in innervated muscle. To assess the functional role of rER in Ca^2+^ regulation at the endplate, we evaluated the response of live isolated FDB fibers to m-3M3-FBS, a potent activator of phospholipase C (PLC) that induces the production of IP3 and leads to an IP3R-dependent Ca^2+^ release in the cytosol. The amplitude of the Ca^2+^ peak induced by m-3M3-FBS treatment was stronger in the sub-synaptic region of the myofiber, compared to non-synaptic region (Supplementary Fig. S4B, C). This was consistent with the strong expression of genes encoding IP3Rs in sub-synaptic nuclei (see Supplementary Fig. S1D) and the accumulation of IP3R1 at the endplate ^48^. The specificity of IP3R-dependent response to m-3M3-FBS was validated by the absence of Ca^2+^ peak in the presence of Xestospongin C, a specific IP3R inhibitor (Supplementary Fig. S4B). These results confirm that rER enrichment at the endplate provides an easily mobilizable Ca^2+^ store that may underlie local SOCE regulation in this region.

We next evaluated the functional consequences of denervation-induced changes in SOCE components expression and localization. To this end, we analyzed SOCE in FDB muscle fibers freshly isolated from innervated muscle or 12 hours and 7 days after denervation (Supplementary Fig. S4D). There was no change in basal cytosolic Ca^2+^ or caffeine-induced Ca^2+^ release in non- and sub-synaptic regions between innervated and denervated muscles (Supplementary Fig. S4E, F). Interestingly, the amplitude of SOCE induced in non- and sub-synaptic regions increased after denervation (Fig. 4G, H). Moreover, SOCE-dependent Ca^2+^ accumulation increased 12 hours after nerve injury in non-synaptic regions, and after 7 days in the sub-synaptic region, compared to innervated muscle (Supplementary Fig. S4G, H). We thus proposed that higher STIM2 expression, sarcotubular remodeling and STIM1 re-localization at the endplate enhance SOCE after denervation in non- and sub-synaptic regions of the myofiber. Together, these results indicate that nerve injury triggers regionalized changes in SOCE, which supports the hypothesis of a role in sensing innervation and initiating the muscle response to denervation.

### Loss of STIM1, but not of STIM2, impairs denervation-induced synaptic gene up-regulation

Given SOCE regionalization in muscle fiber and the changes induced by denervation, we examined whether SOCE in sub- and/or non-synaptic regions contributes to the muscle response to nerve injury. To this end, we injected AAV-short hairpin RNA (shRNA) directed against *Stim1/2* in TA muscle, and 4 weeks later, one limb was denervated for 3 days to evaluate muscle response to denervation (Fig. 5A). A single injection of AAV-sh*Stim1,* AAV-sh*Stim2,* or AAV-sh*Stim1* and AAV-sh*Stim2,* was sufficient to reduce STIM1 and/or STIM2 expression in TA muscle at both mRNA (Supplementary Fig. S5A, B) and protein (Fig. 5B) levels, as compared to muscle injected with AAV-shScramble (control muscle). Knocking down *Stim1* and/or *Stim2* did not perturb body or muscle masses (Supplementary Fig. S5C, D) and did not alter muscle histology (Supplementary Fig. S5E). We next examined the impact of *Stim1/2* knockdown on synaptic gene expression in innervated and denervated muscles. In innervated conditions, repression of synaptic genes relies on the inhibition of *Myog* expression by the combined action of transcriptional repressors, including DACH2 and MITR, an alternative spliced isoform of HDAC9 ^7, 16^. After nerve injury, HDAC4 induction leads to *Dach2* and *Hdac9* repression, which releases the expression of *Myog* and in turn of synaptic genes (Fig. 5C). Knocking down *Stim1* and/or *Stim2* in innervated muscle had limited effects on synaptic gene expression, with only an increased expression of *Chrng* upon *Stim1* down-regulation, as compared to control muscle (Fig. 5D-G). Interestingly, *Stim1* knockdown impinged denervation-induced up-regulation of *Hdac4* (Fig. 5D), *Myog* (Fig. 5E), *Chrna1* (Fig. 5F), and *Chrng* (Fig. 5G) expression 3 days after nerve injury. Other targets of HDAC4, such as *Mitr, Dach2* and *Eno3* were not differentially regulated upon *Stim1* down-regulation (Supplementary Fig. S5F-H). *Stim2* and *Stim1/Stim2* knockdown had no major effect on synaptic gene expression, as compared to control muscle (Fig. 5E). However, their knockdown led to higher expression of *Hdac4*, *Myog, Chrna1* and/or *Chrng,* compared to muscle injected with AAV-sh*Stim1* (Fig. 5D-G). Lastly, to determine if changes in synaptic gene regulation detected upon STIM1/2 depletion alter endplate innervation and morphology, we stained isolated EDL fibers with BTX and antibodies against neurofilament and synaptophysin, to visualize the post- and pre-synaptic NMJ compartments (Supplementary Fig. S5I). Downregulation of *Stim1, Stim2* or both altered neither endplate innervation (Fig. 5H), nor its fragmentation (Fig. 5I) in innervated muscle. In these conditions, endplate fragmentation remained unchanged after a short period (3-days) of denervation (Fig. 5I). Together, these results indicate that STIM1-dependent SOCE plays a key role in sensing denervation, initiating the muscle response likely in the sub-synaptic region, and inducing synaptic gene up-regulation.

**Fig. 5.**
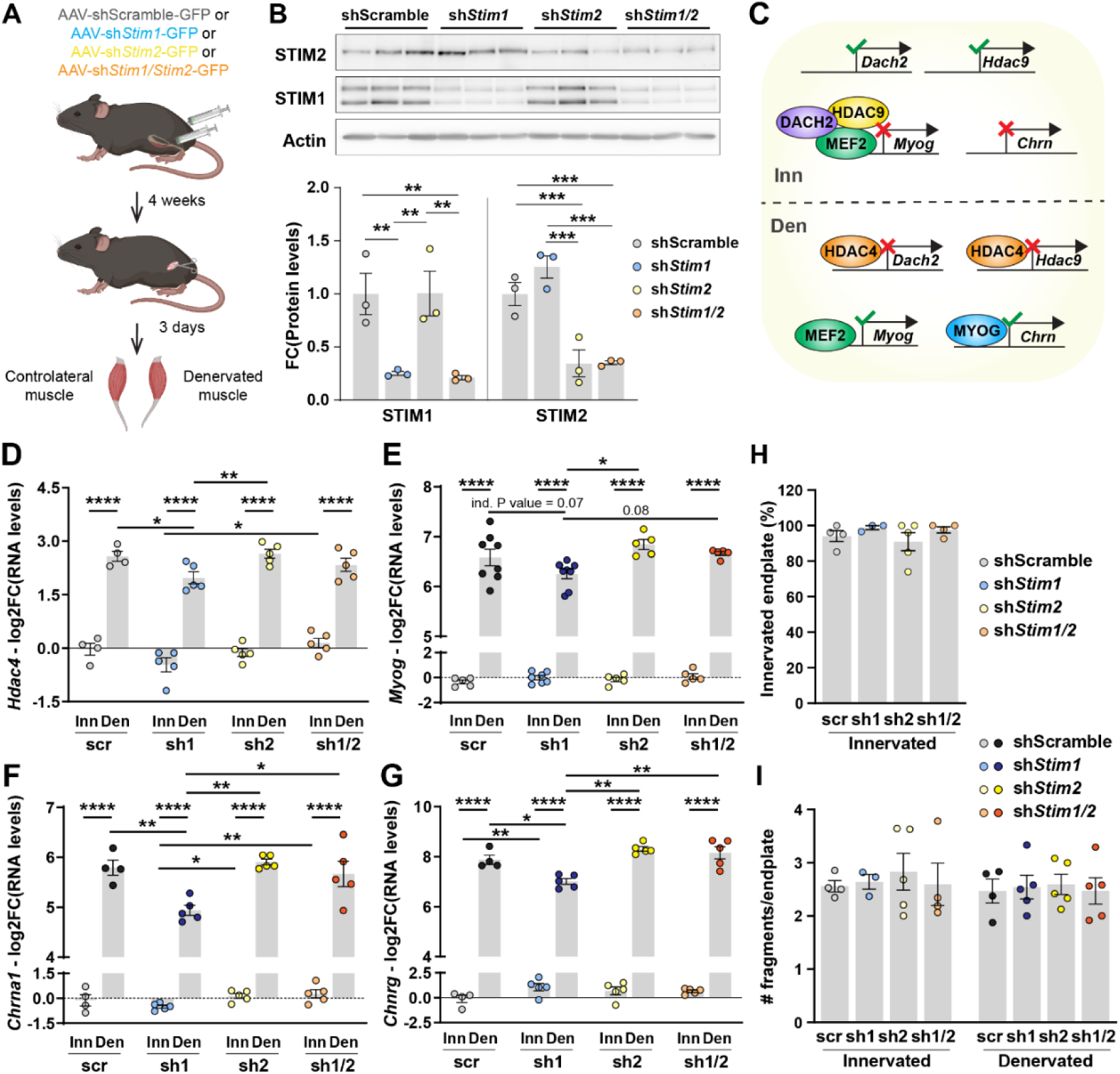
*Stim1* knockdown alters muscle response to denervation. (**A**) Workflow for AAV-based knockdown strategy to evaluate the role of STIM1 and STIM2 in the regulation of synaptic genes in innervated and denervated muscles. (B) Immunoblots of STIM1 and STIM2 in innervated TA muscles infected with AAV-sh-Scramble, -sh*Stim1*, -sh*Stim2*, or –sh*Stim1/Stim2*. Protein levels are normalized on Actin and relative to shScramble control innervated. n=3. (**C**) Schematic representation of the pathway regulating synaptic gene expression in innervated and denervated muscles. (**D-G**) mRNA levels of *Hdac4* (**D**), *Myog* (**E**), *Chrna1* (**F**) and *Chrng* (**G**) in innervated and denervated (3 days) TA muscles infected with AAV-sh-Scramble (scr), -sh*Stim1* (sh1), -sh*Stim2* (sh2), or –sh*Stim1/Stim2* (sh1/2). RNA levels are normalized to *Tbp* and expressed as log2(fold change) of control innervated muscle injected with AAV-shScramble. n=4 for scr (except for E, n=5Inn, 8Den), 5 for sh1 (except for E, n=7Inn, 8Den), sh2, sh1/2 (D-G). (**H**) Proportion of innervated endplates in EDL muscle infected with AAV-shScramble (scr), -sh*Stim1* (sh1), -sh*Stim2* (sh2), or –sh*Stim1/Stim2* (sh1/2), based on the colocalization of neurofilament and synaptophysin immunostaining, with BTX staining. n=4scr, 3sh1, 5sh2, 4sh1/2. (**I**) Number of fragments per endplate in innervated and denervated (3 days) EDL muscles infected with AAV-shScramble (scr), -sh*Stim1* (sh1), -sh*Stim2* (sh2), or –sh*Stim1/Stim2* (sh1/2). n=4scr, 3sh1, 5sh2, 4sh1/2 (except for Den, sh1 and sh1/2, n=5). All values are mean ± s.e.m; *P < 0.05, **P < 0.01, ***P < 0.001, ****P < 0.0001; one-way ANOVA (B) or two-way ANOVA (D-G) with Benjamini and Hochberg’s post-hoc analysis to correct for multiple comparisons. Ind.: individual P value.

### STIM1 overexpression mimics the denervation effect

Given the consequences of *Stim1* knockdown in innervated and denervated muscles, we next evaluated the effect of the overexpression of the short, ubiquitous isoform of STIM1 on the muscle response to denervation. To this end, we injected AAV-STIM1 or AAV-eGFP (control) in TA muscle, and 3 weeks later, one limb was denervated for 3 days (Fig. 6A). A single injection of AAV-STIM1 increased STIM1 expression at both mRNA (Supplementary Fig. S6A) and protein (Fig. 6B) levels, as compared to control muscle. STIM1 overexpression did not lead to compensatory changes in *Stim2* mRNA (Supplementary Fig. S6B) or protein (Fig. 6B) levels. Moreover, there was no change in muscle histology (Supplementary Fig. S6C) upon STIM1 overexpression. We next evaluated the consequences of STIM1 overexpression on synaptic gene expression in innervated and 3-days-denervated muscles. Strikingly, STIM1 overexpression was sufficient to induce a strong increase in *Myog* (Fig. 6C), *Chrna1* (Fig. 6D) and *Chrng* (Fig. 6E) expression in innervated muscle, as compared to control. mRNA levels remained, however, lower as compared to the levels reached after denervation. Moreover, up-regulation of *Myog* and synaptic genes occurred independently from the HDAC4 pathways, as there was no concomitant change in *Hdac4* (Fig. 6F), *Dach2* (Supplementary Fig. S6D), *Mitr* (Supplementary Fig. S6E) and *Eno3* (Supplementary Fig. S6F) expression in innervated muscle upon STIM1 overexpression. Surprisingly, overexpressing STIM1 reduced the up-regulation of *Hdac4* expression (Fig. 6F) and the repression of *Mitr* and *Eno3* (Supplementary Fig. S6E, F) after denervation. However, RNA levels of *Myog* (Fig. 6C), *Chrna1* (Fig. 6D) and *Chrng* (Fig. 6E) reached similar levels as control denervated muscle. Of note, as observed with *Stim1* knockdown, endplate innervation (Fig. 6G) and endplate fragmentation (Fig. 6H and Supplementary Fig. S6G) were unchanged after STIM1 overexpression. These results confirm that STIM1 is involved in the muscle response to denervation and that finely tuned SOCE is required for synaptic genes regulation.

**Fig. 6.**
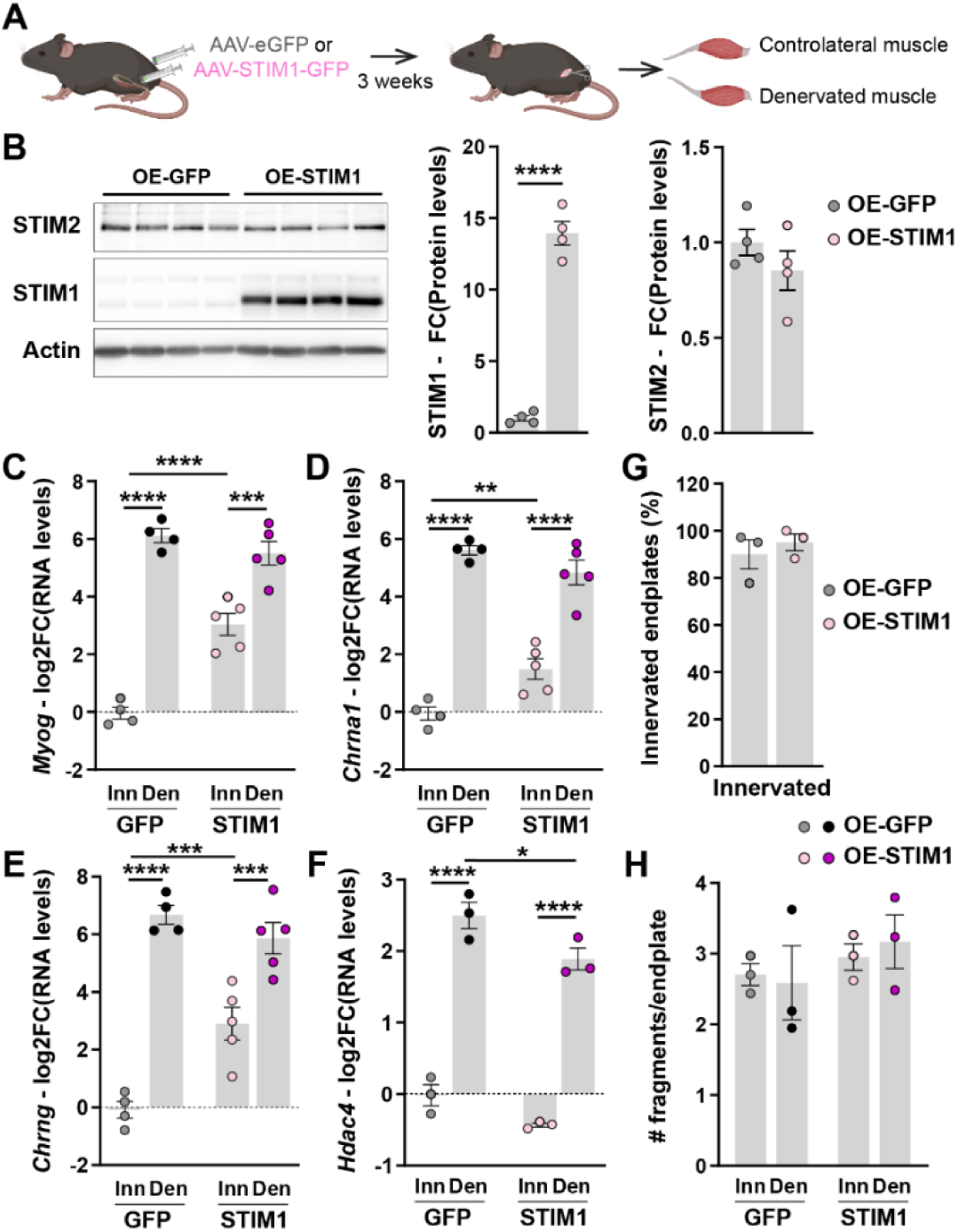
STIM1 overexpression impairs muscle response to denervation. (**A**) Workflow for AAV-based overexpression strategy to evaluate the role of STIM1 in sensing muscle innervation/denervation. (**B**) Immunoblots of STIM1 and STIM2 in innervated TA muscles infected with AAV-eGFP or -STIM1. Protein levels are normalized to Actin and relative to control innervated muscle injected with AAV-eGFP. n= 4. (**C-F)** mRNA levels of *Myog* (**C**), *Chrna1* (**D**), *Chrng* (**E**) and *Hdac4* (**F**) in innervated and denervated (3 days) TA muscles infected with AAV-eGFP or -STIM1. Levels are normalized on *Tbp* and expressed as log2(fold change) of control innervated muscle injected with AAV-eGFP. n=4Inn/5Den (except for F, n=3). (**G**) Proportion of innervated endplates based on the colocalization of Neurofilament and Synaptophysin immunostaining, with BTX staining of innervated EDL bundles. n= 3. (**H**) Number of fragments per endplate in innervated and denervated (3 days) EDL muscles infected with AAV-eGFP or -STIM1. n=3. All values are mean ± s.e.m; **P < 0.01, ***P < 0.001, ****P < 0.0001; unpaired two-tailed Student’s t-test (B), two-way ANOVA with Benjamini and Hochberg’s post-hoc analysis to correct for multiple comparisons (C-F).

### mTORC1 enhances STIM1-dependent SOCE in muscle fibers

Previous reports identify the mTORC1 signaling pathway as a key regulator of NMJ physiology: mTORC1 is activated in response to denervation and it drives NMJ structural instability during aging ^11, 30, 49, 50^. To assess potential functional interactions between SOCE and mTORC1, we first investigated whether mTORC1 modulation affects SOCE. To this end, we used TSCmKO mice, which are depleted for the mTORC1 inhibitor, TSC1, in skeletal muscle, and therefore characterized by constant mTORC1 activation ^51^. The expression of SOCE components was analyzed in innervated and 7-days-denervated muscles from 3-month-old control and TSCmKO mice, *i.e.,* when mutant mice do not show muscle alterations ^52^. While the expression of most SOCE components was unchanged in TSCmKO muscle, STIM1 protein levels were higher in innervated and denervated TA muscles from TSCmKO mice, compared to control (Fig. 7A and Supplementary Fig. S7A). Interestingly, immunostaining for STIM1 of muscle fibers isolated from TSCmKO EDL muscles showed STIM1 mislocalization at the endplate in innervated fibers, which was reminiscent of the effect of denervation observed in control mice (Fig. 7B). The proportion of fibers with a diffuse STIM1 staining further increased in TSCmKO muscle 7 days after nerve injury, as compared to denervated control mice or innervated mutant muscle (Fig. 7C). To evaluate the functional consequences of these changes on SOCE, we measured Ca^2+^ fluxes in FDB fibers isolated from control and TSCmKO mice. Interestingly, SOCE induced after TG and caffeine treatments was higher in sub- and non-synaptic regions of innervated fibers from TSCmKO mice, compared to control mice (Fig. 7D, E). Moreover, there was no difference in SOCE amplitude between non- and sub-synaptic regions in mutant fibers (Fig. 7E), and SOCE did not further increase upon denervation in TSCmKO muscle (Fig. 7F). To determine if these changes are associated with ultrastructural modifications in TSCmKO muscle, we performed EM on innervated and denervated (12 hours and 7 days) EDL muscles. Interestingly, innervated TSCmKO fibers showed dilated T-tubules, as observed after 12 hours of denervation in control mice (Fig. 7G, H, and Supplementary Fig. S7B, C), as well as “pentameric” forms of triads (Fig. 7G), which were previously associated with denervation ^41, 53–56^. In contrast to control muscle, T-tubule dilatation did not further increase in TSCmKO fibers 12 hours after denervation (Fig. 7G, H). Similarly, the surface of SR terminal cisternae was overall higher in TSCmKO muscle compared to control, and it did not differ between innervated and 12-hours-denervated muscles (Fig. 7I). Of note, this effect was mild as compared to the effect of 7 days of denervation observed in control mice. In the sub-synaptic region of innervated mutant fibers, we observed cytosolic spaces enriched in rER, free ribosomes and mitochondria, which were further enlarged after 12 hours of denervation. Notably, mitochondria were smaller and less dense in cristae, and established a high number of contact points with the rER. These changes may also contribute to abnormal Ca^2+^ handling in TSCmKO muscle. Finally, after 7 days of denervation, many mutant fibers showed major ultrastructural alterations, with misaligned and sometimes degenerated sarcomeres associated with cytosolic space, both in non- and sub-synaptic regions (Fig. 7G and Supplementary Fig. S7B). Together, these results indicate that constant mTORC1 activation in skeletal muscle triggers higher SOCE and ultrastructural changes, similar to denervation.

**Fig. 7.**
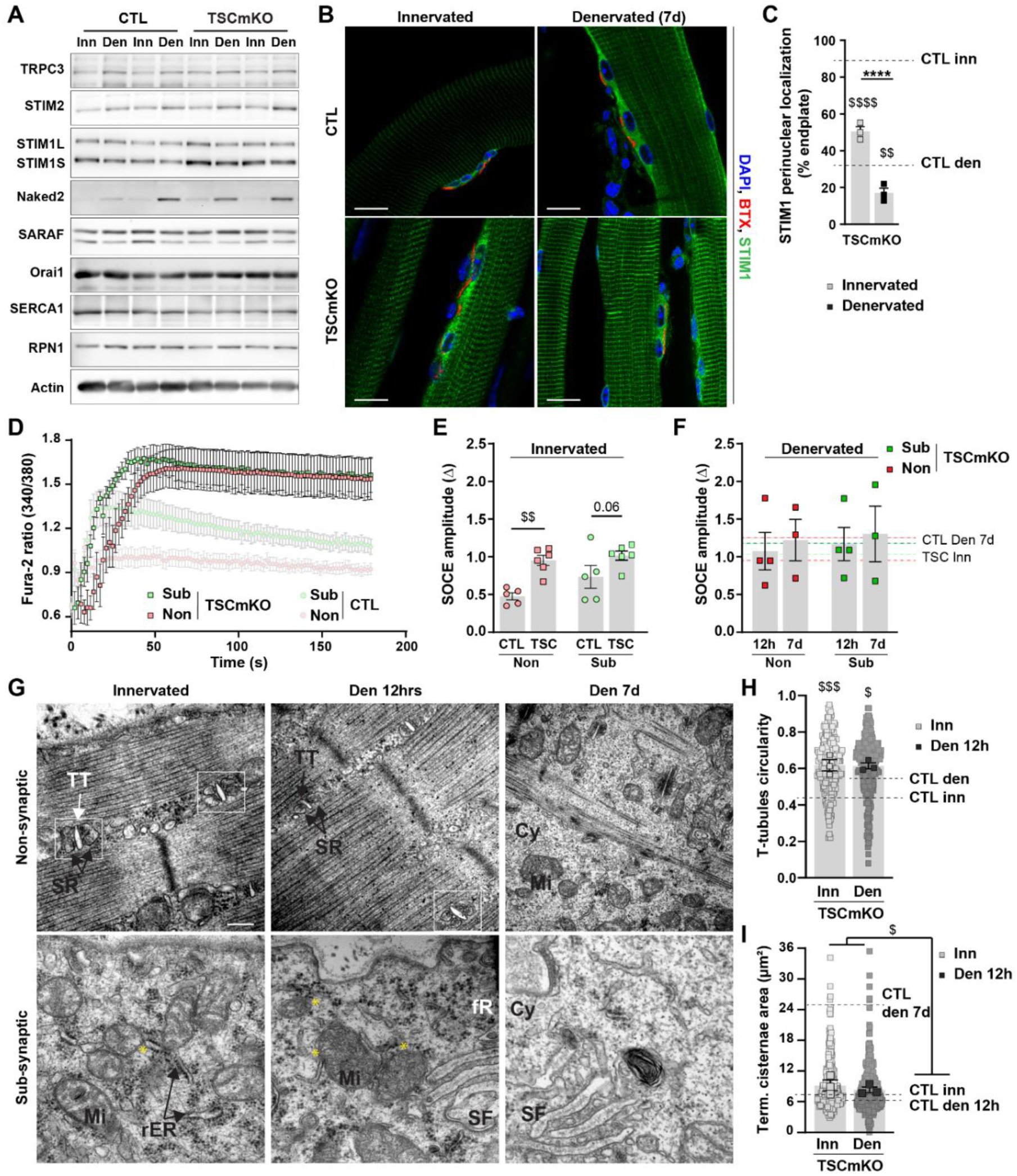
mTORC1 constant activation mimics denervation-induced increase in SOCE capacity and reticulum remodeling. (**A**) Representative immunoblots of proteins involved in SOCE in innervated (Inn) and denervated (Den, 7 days) TA muscles from control (CTL) and TSCmKO mice. Protein levels, relative to Actin, are given in Supplementary Material, Fig. S7A. (**B, C**) Confocal images of STIM1-Cter immunostaining on innervated and denervated (7 days) EDL muscle fibers isolated from control (CTL) and TSCmKO mice. α-Bungarotoxin (BTX) is used to visualize the sub-synaptic region. Scale bar, 20 µm. The proportion of fibers with STIM1 perinuclear localization in the sub-synaptic region is given in **C**. n=3. (**D**) Mean SOCE traces at non- and sub-synaptic regions of fibers isolated from TSCmKO FDB muscle. Light SOCE traces correspond to control muscle. (**E, F**) SOCE amplitude in non- and sub-synaptic regions of innervated (**E**) and denervated (**F**, 12hrs and 7 days) FDB muscle fibers from TSCmKO and control mice. Dashed lines in **F** indicate mean values of control denervated (CTL) and TSCmKO innervated (TSC) muscles. n=5CTL/6TSC (**E**), 4 Den, 12hrs and 3 Den, 7days (**F**). (**G**) Representative images by transmission electron microscopy of longitudinal sections of innervated and denervated (Den, 12 hrs or 7 days) EDL muscles from TSCmKO mice, in non- and sub-synaptic regions. Selections indicate pentameric triads. TT, Mi, SR, SF, rER, fR, Cy indicate T-tubules, mitochondria, sarcoplasmic reticulum, sarcoplasmic folds, rough endoplasmic reticulum, free ribosomes and cytoplasmic space, respectively. Yellow asterisks point to rER/mitochondria contact points. Scale bar, 200 nm. (**H**) T-tubules circularity in innervated (Inn) and denervated (Den, 12 hrs) EDL muscles from TSCmKO mice, quantified based on electron micrographs. Light and opaque dots represents a T-tubule and the mean for one mouse, respectively. Dashed lines indicate mean values of control mice (CTL). n=3. (**I**) SR terminal cisternae surface in innervated (Inn) and denervated (Den, 12 hrs) EDL muscles from TSCmKO mice, quantified based on electron micrographs. Light and opaque dots represents a SR terminal cisternae and the mean for one mouse, respectively. Dashed lines indicate mean values of control mice (CTL). n=3. All values are mean ± s.e.m; ****P < 0.0001 between conditions (Inn/Den); ^$^P < 0.05, ^$$^P < 0.01, ^$$$^P < 0.001, ^$$$$^P < 0.0001 between genotypes; two-way ANOVA with Benjamini and Hochberg’s post-hoc analysis to correct for multiple comparisons (C, E, H, I).

## Discussion

Previous reports establish a role for Ca^2+^ in the spatiotemporal regulation of synaptic gene expression and synaptic protein dynamics, associated with the establishment and maintenance of functional NMJs in skeletal muscle ^2-4,8,9^. However, the molecular mechanisms underlying the cellular regionalization of these Ca^2+^-dependent processes remain largely unknown. Here, we identify a functional regionalization of SOCE in muscle fibers that may lead to the establishment of distinct Ca^2+^ regulatory micro-domains in sub- and non-synaptic regions (Fig. 8). Consistent with a role in maintaining neuromuscular integrity, disruption or exacerbation of SOCE in muscle fibers altered synaptic gene expression and the muscle response to denervation.

**Fig. 8.**
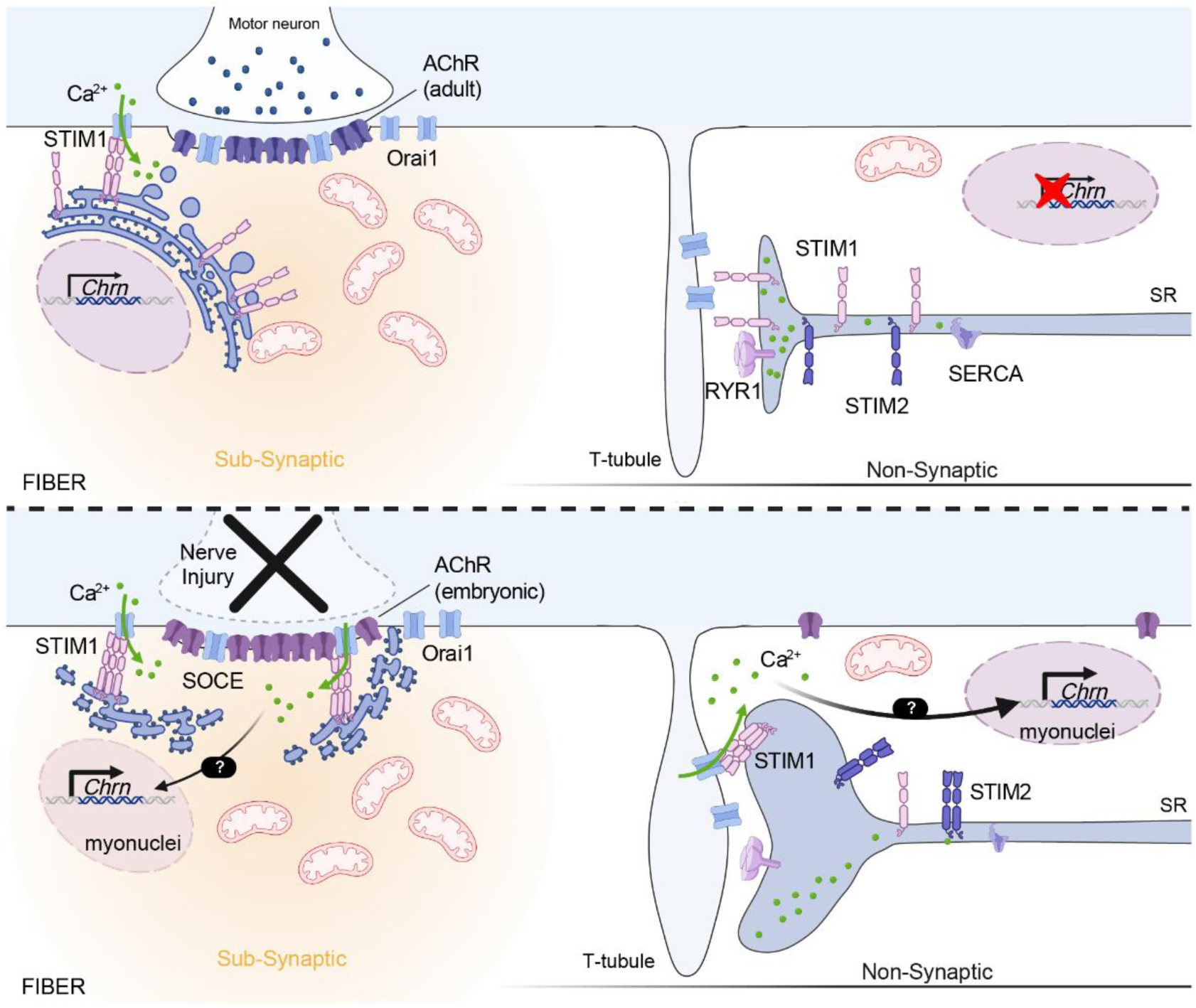
Model depicting the role of SOCE in synaptic gene regulation in non- and sub-synaptic muscle regions. Innervated and denervated muscle fibers are depicted on top and bottom, respectively. SOCE-associated Ca^2+^ fluxes are shown with green arrows. SR: sarcoplasmic reticulum.

Data from the literature suggest that SOCE is involved in the maintenance of SR Ca^2+^ stores in muscle fibers, thereby limiting muscle fatigue. However, the amount of Ca^2+^ mobilized by SOCE has been shown to be relatively small as compared to intracellular stores and to the amount of Ca^2+^ released upon excitation/contraction, interrogating on the role of these fluxes in muscle physiology ^24–29, 57^. Here, we provide a new signaling role for SOCE in myofibers. Indeed, we show that expression of the main components of SOCE and of some regulatory proteins is regionalized in muscle fibers: STIM1, Orai1 and SARAF are enriched in the sub-synaptic region of muscle fibers, while STIM2 is more expressed in non-synaptic regions. Consistently, we identify SOCE in response to rER/SR Ca^2+^ depletion in both sub- and non-synaptic regions of isolated muscle fibers, with higher induced Ca^2+^ accumulation at the endplate. This suggests that SOCE can generate Ca^2+^ micro-domains that differ between non- and sub-synaptic regions and thereby induce distinct regulatory signaling pathways within the muscle fiber (Fig. 8). One can hypothesize that regionalized SOCE reflects the enrichment in rER at the endplate, which is easier to empty than the large SR Ca^2+^ stores present in non-synaptic regions. Ca^2+^ leak from SR is also lower as compared to rER ^26, 58^. The presence of the two different types of reticulum in muscle fibers may contribute to the biphasic pattern (*i.e.,* slow and fast) of SOCE in skeletal muscle, as opposed to monophasic SOCE in undifferentiated cells ^59^. Importantly, we establish that SOCE increased in non- and sub-synaptic regions in denervated muscle fibers, which supports the idea that SOCE responds to nerve injury and triggers specific signaling cascades. SOCE in sub- and non-synaptic regions may contribute to the local regulation of activity-dependent processes, such as synaptic gene regulation and/or synaptic protein turnover (Fig. 8). Consistently, acute *Stim1* knockdown had no major effect on the expression of synaptic genes in innervated muscle, but reduced their up-regulation after denervation. Inversely, acute STIM1 overexpression was sufficient to increase synaptic gene expression in innervated muscle. Thus, increase in STIM1-dependent SOCE after nerve injury may constitute an important signaling event in the release of *Myog* and synaptic gene repression. The class II HDACs, HDAC4/5 and MITR, have opposite effects on synaptic gene expression but are both regulated by CaMKs ^60^. Changes in their activity may hence mediate some of the effects of STIM1 modulation. While STIM1 overexpression increases *Myog* and *Chrna1/g* expression in innervated muscle, it did not perturb the expression of other HDAC4 targets (*e.g. Dach2, Mitr*). Similarly, *Stim1* knockdown limited synaptic gene up-regulation upon denervation, without hampering HDAC4-dependent repression of *Dach2* and *Mitr*. STIM1 may hence promote synaptic gene expression by inhibiting the activity of Ca^2+^-dependent repressors of *Myog* (*e.g.* MITR), rather than promoting HDAC4 activity. Alternatively, SOCE may induce locally the calcineurin-NFAT signaling pathway, previously involved in myogenin regulation ^61–63^. As *Stim2* knockdown normalized the effect of *Stim1* knockdown, STIM2 may exert an opposite effect as compared to the permissive effect of STIM1, by rather inhibiting HDAC4 or promoting MITR activity. Of note, STIM1 overexpression slightly reduced the expression of *Hdac4* and the repression of *Mitr* in denervated muscle, suggesting that excessive STIM1 induction may also end up to inhibit HDAC4 activity.

The mechanisms underlying denervation-induced increase in non-synaptic SOCE remain unclear. Indeed, SOCE is activated in response to a decrease in rER/SR Ca^2+^ levels. The events triggering a potential Ca^2+^ depletion in response to denervation remains to be identified. One intriguing observation was the transient dilatation of T-tubule in muscle fibers detected as early as 12 hours after denervation. This is reminiscent of the transient morphological adaptation of T-tubules to heavy load resistance exercise in human muscle fibers ^64^. These adaptions were attributed to an accumulation of Ca^2+^ in longitudinal T-tubules interacting with sarcomeres, to avoid Ca^2+^ accumulation in cytosol and associated fiber damage. Our observations suggest that similar adaptations occur in muscle response to denervation to help maintain fiber viability. Whether this cyto-architectural change triggers the fast increase in SOCE capacity in non-synaptic region after denervation remains to be evaluated. Similarly, the mechanisms involved in SOCE regulation in the sub-synaptic region are unknown. The enrichment of STIM1 at the endplate corroborates a recent study showing that STIM1 physically interacts with desmin ^36^, a protein known to be enriched in NMJ post-synaptic compartment ^65^. Given that desmin is degraded in denervated muscle ^66^, the interaction with STIM1 may be lost and drive STIM1 re-localization. Altered SOCE regulation may thus contribute to the dysregulation of synaptic gene expression and NMJ dysfunction reported in desmin-deficient mice ^67^. Importantly, we showed that STIM1/Orai1 enrichment and high SOCE capacity in the sub-synaptic region are associated with clusters of mitochondria present between the endplate membrane folds and sub-synaptic nuclei. Given the functional link between SOCE and mitochondrial Ca^2+^ homeostasis, sub-synaptic mitochondria may be involved in SOCE-dependent regulation of synaptic gene expression and of the muscle response to denervation. In particular, this could explain why mitochondria dysfunction is sufficient to dismantle NMJs ^68–70^. Of note, a recent study demonstrated that the inducible loss of STIM1 in adult fibers is associated with important changes in fiber metabolism ^71^. In this context, SOCE has been proposed to act as a rheostat to fine-tune mitochondrial function to muscle contraction. Our results reinforce the proposed role for STIM1 as a sensor of muscle activity with an additional signaling role to sustain synaptic gene expression.

While the mTORC1 pathway has been shown to play a key role in NMJ maintenance and the muscle response to denervation, downstream targets mediating this effect are not well understood ^11, 30, 50, 72^. We identify that constant mTORC1 activation in TSCmKO mice results in a strong increase in SOCE capacity. Consistently, mTORC1 activation increases STIM1 expression and SOCE capacity in cancer cells ^73^. Inversely, SOCE induces the Akt/mTORC1 pathways during osteogenesis, as well as in cardiomyocytes ^74, 75^ indicating that these two signaling pathways are interconnected. The reported induction of mTORC1 in response to denervation ^30^ may hence trigger or alternatively rely on the increase in SOCE in denervated muscle. The mechanisms underlying the effect of mTORC1 on STIM1 expression and SOCE capacity remain unclear. TSCmKO mice showed T-tubule dilation in innervated muscle that is reminiscent of what we observed 12 hours after denervation in control mice, when non-synaptic SOCE capacity is increased. mTORC1-dependent ultrastructural changes may hence support higher SOCE. ER stress, reported in TSCmKO skeletal muscle ^76^, has been associated with higher SOCE capacity in other organs ^77^ and may thus contribute as well, to mTORC1-dependent increase in SOCE. Chronic increase in SOCE-associated signaling may contribute to the defective response of TSCmKO muscle to nerve injury by blunting denervation sensing and signaling cascade induction, in contrast to STIM1 acute overexpression. Several reports have highlighted the central role of mTORC1 dysregulation in NMJ integrity loss upon aging ^11, 50, 72, 78^. Importantly, changes in SOCE have also been linked to aging-associated muscle dysfunction ^79, 80^. Whether these changes alter NMJ stability and occur upstream or downstream of mTORC1 dysregulation is not known. Similarly, gain-of-function mutations in *STIM1* cause tubular aggregate myopathies mainly characterized by muscle weakness and cramps ^20^. To date, there was no report of NMJ dysfunction in these patients. However, the results obtained with STIM1 overexpression suggest that *STIM1* gain-of-function mutations may alter synaptic gene expression and progressively impair the maintenance of NMJs. Identifying new pathomechanisms involved in muscle dysfunction in these myopathies would open new directions for the development of treatments for these diseases.

In conclusion, our work has unraveled a new role for SOCE in muscle fibers, as an important regulator of activity-dependent regionalized expression of synaptic genes and sensor of muscle denervation. SOCE inhibition may represent a new strategy to mitigate neuromuscular affliction in pathological conditions associated with mTORC1 activation, such as sarcopenia.

## Methods

### Animals

Generation and genotyping of TSCmKO transgenic mice were described previously ^52^. Control mice for TSCmKO mice were littermates that were floxed for *Tsc1* but did not express Cre-recombinase. For muscle denervation, sciatic nerve was cut unilaterally on mice anesthetized with isoflurane, as previously described^51^. The contralateral leg serves as control. For AAV study, anterior hindlimb compartments of 2/3-months-old C57BL6/J mice were injected with adeno-associated virus serotype 9 (AAV9 – Vector Biolabs) carrying either transgenes for STIM1 (AAV9-CMV-mStim1-2A-eGFP) or GFP (AAV9-CMV-eGFP), or shRNA directed against for *Stim1* (AAV9-GFP-U6-*mStim1*-shRNA), or *Stim2* (AAV9-GFP-U6-*mStim2*-shRNA) or a scrambled control (AAV9-GFP-U6-scramble-shRNA) at a dose of 8 × 10^10^ viral particles. All mouse experiments were in accordance with the European Union guidelines on animal experimentation and approved by the Veterinary Office of the Canton of Geneva (application number GE220/GE227).

### Transcript expression analyses

Total RNA was extracted with the RNeasy Fibrous Tissue Mini Kit (Qiagen). Quantitative PCR was performed on DNAse-treated RNA, reverse transcribed with the High-Capacity cDNA Reverse Transcription Kit (Applied Biosystems), amplified with the Power UP Sybr Green Master Mix (Applied Biosystem). Data were analyzed using the StepOne software and normalized to *Tbp* expression. Primers are listed in Supplementary Table 1.

### Western blotting

Nitrogen-powdered TA muscles were lysed in RIPA buffer (50 mM Tris HCl pH8, 150 mM NaCl, 1% NP-40, 0.5% sodium deoxycholate, 0.1% SDS, 1% Triton-X, 10% glycerol) with protease (Pierce) and phosphatase (Roche) inhibitor cocktail tablets. Lysates were incubated at 4°C on rotating wheel for 2 hrs, sonicated two times for 10 s and centrifuged at 10,000 rpm for 20 min at 4 °C. Cleared lysates were used to determine total protein amount (BCA Protein Assay, Pierce). Proteins were separated in polyacrylamide SDS gels and transferred to nitrocellulose membrane (Amersham). Membranes were blocked in 3% bovine serum albumin (BSA) in tris-buffered saline with 0.1% Tween-20, incubated successively with primary and appropriate HRP-conjugated secondary antibodies, and revealed using the LumiGLO® Peroxidase Chemiluminescent Substrate Kit (SeraCare). Light emission was recorded using a chemiluminescent detection system (iBright, ThermoFischer scientific) and quantified using Fiji software [v1.53, National Institutes of Health (NIH)].

### Immunostaining and inorganic staining on muscle sections

Muscles were frozen in liquid nitrogen-cooled isopentane; eight µm muscle cryosections were used for histology analyses. Cryostat sections were stained with Hematoxylin/Eosin (HE) according to classical methods ^81^. For immunostaining, sections were fixed with PFA 4% and then blocked with 3% IgG free BSA and AffiniPure Mouse IgG Fab Fragments (Jackson ImmunoResearch) in phosphate-buffered saline (PBS). They were incubated sequentially with primary and secondary fluorescent antibodies, together with α-bungarotoxin-Alexa647 (Invitrogen), and mounted in Vectashield DAPI (Vector). Images were captured using the Zeiss Axio Imager M2 or Axio Imager Z2 Basis LSM 800 microscopes.

### Immunostaining of muscle bundles and single isolated fibers

EDL muscles were fixed on tibia in 2% PFA for 25 min, washed, divided in bundles, and permeabilized in PBS, 1% Triton-X100, 1% BSA. Bundles were then successively incubated with primary antibodies against Neurofilament and Synaptophysin, and the corresponding secondary antibodies (Invitrogen). Alternatively, single fibers were isolated from PFA-fixed EDL muscle, permeabilized with PBS, 5% Triton-X100, 3% BSA and AffiniPure Mouse IgG Fab fragments (Jackson ImmunoResearch), and incubated sequentially with primary and secondary antibodies, with fluorescent α-bungarotoxin to distinguish non- and sub-synaptic regions. Fibers were mounted in Vectashield DAPI (Vector). More than 30 fibers were used per analysis. Images were recorded using the Axio Imager Z2 Basis LSM 800 microscope.

### Antibodies

The following primary antibodies were used: STIM1-Nter (S6072, 1/250 for IF), STIM1-Cter (S6197, 1/250 for IF), Neurofilament 200 (N4142; 1/2000 for IF) from Sigma; Synaptophysin (GT2589; 1/200 for IF) from Genetex; STIM1 (AB9870, 1/250 for IF and 1/1000 for WB) from Millipore; Orai1 (PA5-26378, 1/250 for IF, 1/1000 for WB), TRPC3 (PA5-77307, 1/250 for IF, 1/1000 for WB), SARAF (PA5-24237, 1/250 for IF, 1/1000 for WB) from Invitrogen; Naked2 (#2073, 1/250 for IF, 1/500 for WB), STIM2 (#4917, 1/250 for IF, 1/500 for WB), Pan-CaMKII-P (#12716, 1/250 for IF, 1/1000 for WB), Pan-CaMKII (#4436, 1/250 for IF, 1/500 for WB) from Cell Signaling; SERCA1 (CaF2-5D2, 1/100 for IF, 1/250 for WB) from DSHB; β-Actin (47778, 1/10 000 for WB) from Santa Cruz. RPN1 (1/1000 for WB) and KDEL (1/250 for IF) antibodies were gift from the PHYM department (University of Geneva).

### Transmission electron microscopy

EDL muscles were fixed with 2.5% glutaraldehyde in 0.1 M phosphate buffer, pH 7.2, washed with phosphate buffer, treated with 2% osmium tetraoxyde and immersed in 0.25% uranyl acetate over night to enhance contrast of membranes. Small EDL pieces were gradually dehydrated in ethanol, substituted in a mix of propylene oxyde-epon and embedded in Epon (Delta microscopy). Thin sections (70 nm) were collected onto 200 mesh copper grids, and counterstained with uranyl acetate and lead citrate. Grids were examined with a Morgagni electron microscope (FEI Company, Eindhoven, Netherlands) operating at 80 kV. Images were acquired with a charge-coupled device camera (AMT).

### Ca^2+^ analysis

Single FDB muscle fibers were isolated based on a classical protocol ^82^ that was adapted to minimize the period before analysis. Briefly, for enzymatic digestion, muscles were incubated for 60 min at 37°C with 0.2% collagenase type I (C0130, Sigma-Aldrich) in DMEM (41966, ThermoFisher Scientific) supplemented with BTX-biotin (B1196, Invitrogen). Mechanical disruption was done in DMEM supplemented with streptavidin-Alexa568 (S11226, Invitrogen) using a glass pipet. FDB fibers were transferred into laminin-coated (2 µg/ml, LN211-05, BioLamina) 8 well-slides (80826, ibidi) in Ca^2+^ buffer, loaded with 2 μM of Fura-2 AM (F1221, ThermoFischer Scientific), 0.1 % pluronic acid (Pluronic F-127, P3000MP, ThermoFischer Scientific) and 40 µM of BTS (203895, Sigma-Aldrich) in the dark at room temperature for 60 min, and were washed twice. All solutions used in following steps contained 40 µM of BTS. After 20 min to allow the de-esterification of the dye, fluorescence was recorded using Zeiss Axio Observer A1 microscope (Carl Zeiss AG) equipped with a Lambda DG4 illumination system (Sutter Instrument), which rapidly changed the excitation wavelengths between 340 nm and 380 nm (340AF15 and 380AF15, Omega Optical). Emission was collected through a 415 DRLP dichroic mirror, and a 510WB40 filter (Omega Optical) by a cooled 16-bit CMOS camera (Prime-BSI-Express back-illuminated scientific, Visitron Systems, Puchheim, Germany). Image acquisitions and analysis were performed with the VisiWiew software (Visitron Systems, Puchheim). Cells were stimulated in Ca^2+^-free solution with 1 μM thapsigargin (Tg, T9033, Sigma-Aldrich) for 7 min, 20 mM of caffeine for 3 min, and then 2 mM Ca^2+^ was added to induce SOCE. Alternatively, cells were stimulated in Ca^2+^-free solution with 1 μM Tg and 10 µM dantrolene (251680, Sigma-Aldrich) for 2 min and then with 15 µM phospholipase C activator m-3M3FBS (525185, Sigma-Aldrich) for 2 min. m-3M3FBS action on phospholipase C and IP3R was validated by showing the absence of response with 5µM Xestospongin C (X2628, Sigma-Aldrich). Basal Ca^2+^ corresponds to Ca^2+^ levels before the addition of any drugs. Caffeine-induced Ca^2+^ release is the difference (Δ) between cytosolic Ca^2+^ after and before the addition of caffeine. IP3R-dependent Ca^2+^ is the difference (Δ) between cytosolic Ca^2+^ after and before the addition of m-3M3FBS. SOCE was calculated as the difference (Δ – amplitude) between cytosolic Ca^2+^ after and before the addition of Ca^2+^ in the medium. The area under the curve following Ca^2+^ entry before the peak was analyzed in both non-synaptic and sub-synaptic (area positive for BTX) regions. The bath contained 140mM NaCl, 5mM KCl, 1mM MgCl2, 20mM Hepes, 10mM glucose, and 2mM CaCl2 in Ca^2+^-containing solution, pH 7.4.

### Public data analysis

Analysis of gene expression using public repository data was performed using SarcoAtlas (https://sarcoatlas.scicore.unibas.ch/) ^11^ and Myoatlas (https://research.cchmc.org/myoatlas/) ^33^. We analyzed the ‘NMJ’ data set (GEO accession number: GSE139209) that contains mRNA profiles of synaptic (NMJ) and non-synaptic (xNMJ) regions of TA muscle of 10- and 30-month-old wild type mice.

### Quantification and statistical analysis

Images were analyzed with Fiji software. Sub-synaptic enrichment score is based on the colocalization between signals from immunostaining of the protein of interest and BTX staining that was quantified using the plugin Coloc 2. The Pearson R value was then transformed into t score using the following formula: 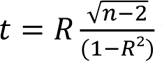 where R is the correlation score (Pearson R) and n the sample size. The P value was calculated from t score using the P-Value calculator (https://www.graphpad.com/quickcalcs/pvalue1.cfm). Circularity and roundness of T-tubules were determined by the shape descriptor plugin of Fiji. Terminal cisternae surface was calculated using the freehand selection tool of Fiji. All data are presented as mean of values of independent samples ± s.e.m. Raw data from each individual experiment were evaluated using an unpaired two-tailed Student’s t-test with 95% confidence. When two experimental conditions were combined, statistical comparisons were done using ordinary two-way ANOVA test. Multiple comparisons were corrected by controlling the false discovery rate (<0.05) using the original method of Benjamini and Hochberg.

## Acknowledgements

The authors thank members of the PFMU, Bioimaging, Histology and ChiRO core facilities (University of Geneva, Switzerland) for experimental support. We thank Prof. M. Rüegg (University of Basel, Switzerland) for the generous gift of TSCmKO mice.

## Author contributions

Conceptualization: AP, PC; Methodology: AP, OD, JR, MF, SK; PC; Investigation: AP, JR, FC; Supervision: AP, PC; Writing original draft: AP, PC; Writing review & editing: AP, PC, SK, MF.

## Data availability

Data generated during the study will be freely accessible on the Yareta repository database upon acceptance.

## References

1 Tintignac, L. A., Brenner, H. R. & Ruegg, M. A. Mechanisms Regulating Neuromuscular Junction Development and Function and Causes of Muscle Wasting. Physiol Rev 95, 809–852 (2015).

2 Kaplan, M. M. et al. Calcium Influx and Release Cooperatively Regulate AChR Patterning and Motor Axon Outgrowth during Neuromuscular Junction Formation. Cell Rep 23, 3891–3904 (2018).

3 Chen, F. et al. Neuromuscular synaptic patterning requires the function of skeletal muscle dihydropyridine receptors. Nat Neurosci 14, 570–577 (2011).

4 Walke, W. et al. Calcium-dependent regulation of rat and chick muscle nicotinic acetylcholine receptor (nAChR) gene expression. J Biol Chem 269, 19447–19456 (1994).

5 Macpherson, P., Kostrominova, T., Tang, H. & Goldman, D. Protein kinase C and calcium/calmodulin-activated protein kinase II (CaMK II) suppress nicotinic acetylcholine receptor gene expression in mammalian muscle. A specific role for CaMK II in activity-dependent gene expression. J Biol Chem 277, 15638–15646 (2002).

6 Tang, H., Sun, Z. & Goldman, D. CaM kinase II-dependent suppression of nicotinic acetylcholine receptor delta-subunit promoter activity. J Biol Chem 276, 26057–26065 (2001).

7. Tang, H. & Goldman, D. Activity-dependent gene regulation in skeletal muscle is mediated by a histone deacetylase (HDAC)-Dach2-myogenin signal transduction cascade. PNAS 103, 16977-16982 (2006).

8 Megeath, L. J. & Fallon, J. R. Intracellular calcium regulates agrin-induced acetylcholine receptor clustering. J Neurosci 18, 672–678 (1998).

9 Martinez-Pena y Valenzuela, I., Mouslim, C. & Akaaboune, M. Calcium/calmodulin kinase II-dependent acetylcholine receptor cycling at the mammalian neuromuscular junction in vivo. J Neurosci 30, 12455–12465 (2010).

10 Koliatsos, V. E., Clatterbuck, R. E., Winslow, J. W., Cayouette, M. H. & Price, D. L. Evidence that brain-derived neurotrophic factor is a trophic factor for motor neurons in vivo. Neuron 10, 359–367 (1993).

11 Ham, D. J. et al. The neuromuscular junction is a focal point of mTORC1 signaling in sarcopenia. Nat Commun 11, 4510 (2020).

12 Aare, S. et al. Failed reinnervation in aging skeletal muscle. Skelet Muscle 6, 29 (2016).

13 Akaaboune, M., Culican, S. M., Turney, S. G. & Lichtman, J. W. Rapid and reversible effects of activity on acetylcholine receptor density at the neuromuscular junction in vivo. Science 286, 503–507 (1999).

14 Yampolsky, P., Pacifici, P. G. & Witzemann, V. Differential muscle-driven synaptic remodeling in the neuromuscular junction after denervation. Eur. J. Neurosci. 31, 646–658 (2010).

15 Brown, M. C. & Ironton, R. Sprouting and regression of neuromuscular synapses in partially denervated mammalian muscles. J Physiol 278, 325–348 (1978).

16 Mejat, A. et al. Histone deacetylase 9 couples neuronal activity to muscle chromatin acetylation and gene expression. Nat. Neurosci. 8, 313–321 (2005).

17 Tang, H. et al. A histone deacetylase 4/myogenin positive feedback loop coordinates denervation-dependent gene induction and suppression. Mol. Biol. Cell 20, 1120–1131 (2009).

18 Cohen, T. J. et al. The histone deacetylase HDAC4 connects neural activity to muscle transcriptional reprogramming. J. Biol. Chem. 282, 33752–33759 (2007).

19 Prakriya, M. & Lewis, R. S. Store-Operated Calcium Channels. Physiol Rev 95, 1383–1436 (2015).

20 Silva-Rojas, R., Laporte, J. & Bohm, J. STIM1/ORAI1 Loss-of-Function and Gain-of-Function Mutations Inversely Impact on SOCE and Calcium Homeostasis and Cause Multi-Systemic Mirror Diseases. Front Physiol 11, 604941 (2020).

21 Protasi, F., Girolami, B., Roccabianca, S. & Rossi, D. Store-operated calcium entry: From physiology to tubular aggregate myopathy. Curr Opin Pharmacol 68, 102347 (2023).

22 Morin, G. et al. Tubular aggregate myopathy and Stormorken syndrome: Mutation spectrum and genotype/phenotype correlation. Hum Mutat 41, 17–37 (2020).

23 Pearce, L. et al. Ryanodine receptor activity and store-operated Ca(2+) entry: Critical regulators of Ca(2+) content and function in skeletal muscle. J Physiol 601, 4183–4202 (2023).

24 Lamboley, C. R. et al. Ryanodine receptor leak triggers fiber Ca(2+) redistribution to preserve force and elevate basal metabolism in skeletal muscle. Sci Adv 7, eabi7166 (2021).

25 Lilliu, E., Koenig, S., Koenig, X. & Frieden, M. Store-Operated Calcium Entry in Skeletal Muscle: What Makes It Different? Cells 10 (2021).

26 Bolanos, P. & Calderon, J. C. Excitation-contraction coupling in mammalian skeletal muscle: Blending old and last-decade research. Front Physiol 13, 989796 (2022).

27 Michelucci, A. et al. Pre-assembled Ca2+ entry units and constitutively active Ca2+ entry in skeletal muscle of calsequestrin-1 knockout mice. J Gen Physiol 152 (2020).

28 Michelucci, A. et al. Transverse tubule remodeling enhances Orai1-dependent Ca(2+) entry in skeletal muscle. Elife 8 (2019).

29 Boncompagni, S., Michelucci, A., Pietrangelo, L., Dirksen, R. T. & Protasi, F. Exercise-dependent formation of new junctions that promote STIM1-Orai1 assembly in skeletal muscle. Sci Rep 7, 14286 (2017).

30 Castets, P. et al. mTORC1 and PKB/Akt control the muscle response to denervation by regulating autophagy and HDAC4. Nat Commun 10, 3187 (2019).

31 Miller, S. G. & Kennedy, M. B. Regulation of brain type II Ca2+/calmodulin-dependent protein kinase by autophosphorylation: a Ca2+-triggered molecular switch. Cell 44, 861–870 (1986).

32 Moresi, V. et al. Myogenin and class II HDACs control neurogenic muscle atrophy by inducing E3 ubiquitin ligases. Cell 143, 35–45 (2010).

33 Petrany, M. J. et al. Single-nucleus RNA-seq identifies transcriptional heterogeneity in multinucleated skeletal myofibers. Nat Commun 11, 6374 (2020).

34 Alkhani, H. et al. Contribution of TRPC3 to store-operated calcium entry and inflammatory transductions in primary nociceptors. Mol Pain 10, 43 (2014).

35 Wu, B. et al. NKD2 mediates stimulation-dependent ORAI1 trafficking to augment Ca(2+) entry in T cells. Cell Rep 36, 109603 (2021).

36. Zhang, H., et al. Desmin interacts with STIM1 and coordinates Ca2+ signaling in skeletal muscle. JCI Insight 6 (2021).

37 Dagan, I. & Palty, R. Regulation of Store-Operated Ca(2+) Entry by SARAF. Cells 10 (2021).

38 Stiber, J. et al. STIM1 signalling controls store-operated calcium entry required for development and contractile function in skeletal muscle. Nat Cell Biol 10, 688–697 (2008).

39 Edwards, J. N. et al. Ultra-rapid activation and deactivation of store-operated Ca(2+) entry in skeletal muscle. Cell Calcium 47, 458–467 (2010).

40 Wei-Lapierre, L., Carrell, E. M., Boncompagni, S., Protasi, F. & Dirksen, R. T. Orai1-dependent calcium entry promotes skeletal muscle growth and limits fatigue. Nat Commun 4, 2805 (2013).

41 Sakakima, H., Kawamata, S., Kai, S., Ozawa, J. & Matsuura, N. Effects of short-term denervation and subsequent reinnervation on motor endplates and the soleus muscle in the rat. Arch Histol Cytol 63, 495–506 (2000).

42 Sigrist, S. J. et al. Postsynaptic translation affects the efficacy and morphology of neuromuscular junctions. Nature 405, 1062–1065 (2000).

43 Launikonis, B. S. et al. Depletion “skraps” and dynamic buffering inside the cellular calcium store. Proc Natl Acad Sci U S A 103, 2982–2987 (2006).

44 Rudolf, R., Magalhaes, P. J. & Pozzan, T. Direct in vivo monitoring of sarcoplasmic reticulum Ca2+ and cytosolic cAMP dynamics in mouse skeletal muscle. J Cell Biol 173, 187–193 (2006).

45 Canato, M. et al. Massive alterations of sarcoplasmic reticulum free calcium in skeletal muscle fibers lacking calsequestrin revealed by a genetically encoded probe. Proc Natl Acad Sci U S A 107, 22326–22331 (2010).

46 Kurebayashi, N. & Ogawa, Y. Depletion of Ca2+ in the sarcoplasmic reticulum stimulates Ca2+ entry into mouse skeletal muscle fibres. J Physiol 533, 185–199 (2001).

47 Koenig, X., Choi, R. H. & Launikonis, B. S. Store-operated Ca(2+) entry is activated by every action potential in skeletal muscle. Commun Biol 1, 31 (2018).

48 Zhu, H., Bhattacharyya, B. J., Lin, H. & Gomez, C. M. Skeletal muscle IP3R1 receptors amplify physiological and pathological synaptic calcium signals. J Neurosci 31, 15269–15283 (2011).

49 Tang, H. et al. mTORC1 underlies age-related muscle fiber damage and loss by inducing oxidative stress and catabolism. Aging Cell 18, e12943 (2019).

50 Ang, S. J. et al. Muscle 4EBP1 activation modifies the structure and function of the neuromuscular junction in mice. Nat Commun 13, 7792 (2022).

51 Bentzinger, C. F. et al. Differential response of skeletal muscles to mTORC1 signaling during atrophy and hypertrophy. Skelet. Muscle 3, 6 (2013).

52 Castets, P. et al. Sustained activation of mTORC1 in skeletal muscle inhibits constitutive and starvation-induced autophagy and causes a severe, late-onset myopathy. Cell Metab 17, 731–744 (2013).

53 Pellegrino, C. & Franzini, C. An Electron Microscope Study of Denervation Atrophy in Red and White Skeletal Muscle Fibers. J Cell Biol 17, 327–349 (1963).

54 Lu, D. X., Huang, S. K. & Carlson, B. M. Electron microscopic study of long-term denervated rat skeletal muscle. Anat Rec 248, 355–365 (1997).

55 Carraro, U. et al. Chronic denervation of rat hemidiaphragm: maintenance of fiber heterogeneity with associated increasing uniformity of myosin isoforms. J Cell Biol 100, 161–174 (1985).

56 Engel, A. G. & Stonnington, H. H. Trophic functions of the neuron. II. Denervation and regulation of muscle. Morphological effects of denervation of muscle. A quantitative ultrastructural study. Ann N Y Acad Sci 228, 68–88 (1974).

57 Rincon, O. A., Milan, A. F., Calderon, J. C. & Giraldo, M. A. Comprehensive Simulation of Ca(2+) Transients in the Continuum of Mouse Skeletal Muscle Fiber Types. Int J Mol Sci 22 (2021).

58 Bygrave, F. L. & Benedetti, A. What is the concentration of calcium ions in the endoplasmic reticulum? Cell Calcium 19, 547–551 (1996).

59 Darbellay, B., Arnaudeau, S., Bader, C. R., Konig, S. & Bernheim, L. STIM1L is a new actin-binding splice variant involved in fast repetitive Ca2+ release. J Cell Biol 194, 335–346 (2011).

60 McKinsey, T. A., Zhang, C. L., Lu, J. & Olson, E. N. Signal-dependent nuclear export of a histone deacetylase regulates muscle differentiation. Nature 408, 106–111 (2000).

61 Kiviluoto, S. et al. STIM1 as a key regulator for Ca2+ homeostasis in skeletal-muscle development and function. Skelet Muscle 1, 16 (2011).

62 Lin, Y. P., Bakowski, D., Mirams, G. R. & Parekh, A. B. Selective recruitment of different Ca(2+)-dependent transcription factors by STIM1-Orai1 channel clusters. Nat Commun 10, 2516 (2019).

63 Kar, P. et al. Dynamic assembly of a membrane signaling complex enables selective activation of NFAT by Orai1. Curr Biol 24, 1361–1368 (2014).

64 Cully, T. R. et al. Human skeletal muscle plasmalemma alters its structure to change its Ca(2+)-handling following heavy-load resistance exercise. Nat Commun 8, 14266 (2017).

65 Askanas, V., Bornemann, A. & Engel, W. K. Immunocytochemical localization of desmin at human neuromuscular junctions. Neurology 40, 949–953 (1990).

66 Aweida, D., Rudesky, I., Volodin, A., Shimko, E. & Cohen, S. GSK3-beta promotes calpain-1-mediated desmin filament depolymerization and myofibril loss in atrophy. J Cell Biol 217, 3698–3714 (2018).

67 Eiber, N. et al. Lack of Desmin in Mice Causes Structural and Functional Disorders of Neuromuscular Junctions. Front Mol Neurosci 13, 567084 (2020).

68 Zhou, J. et al. Hyperactive intracellular calcium signaling associated with localized mitochondrial defects in skeletal muscle of an animal model of amyotrophic lateral sclerosis. J Biol Chem 285, 705–712 (2010).

69 Dupuis, L. et al. Muscle mitochondrial uncoupling dismantles neuromuscular junction and triggers distal degeneration of motor neurons. PLoS One 4, e5390 (2009).

70 Genin, E. C. et al. Mitochondrial defect in muscle precedes neuromuscular junction degeneration and motor neuron death in CHCHD10(S59L/+) mouse. Acta Neuropathol 138, 123–145 (2019).

71 Wilson, R. J. et al. Disruption of STIM1-mediated Ca(2+) sensing and energy metabolism in adult skeletal muscle compromises exercise tolerance, proteostasis, and lean mass. Mol Metab 57, 101429 (2022).

72 Baraldo, M. et al. Skeletal muscle mTORC1 regulates neuromuscular junction stability. J Cachexia Sarcopenia Muscle 11, 208–225 (2020).

73 Peng, H. et al. mTORC1 enhancement of STIM1-mediated store-operated Ca2+ entry constrains tuberous sclerosis complex-related tumor development. Oncogene 32, 4702–4711 (2013).

74 Huang, Y., Li, Q., Feng, Z. & Zheng, L. STIM1 controls calcineurin/Akt/mTOR/NFATC2-mediated osteoclastogenesis induced by RANKL/M-CSF. Exp Ther Med 20, 736–747 (2020).

75 Bonilla, I. M. et al. STIM1 ablation impairs exercise-induced physiological cardiac hypertrophy and dysregulates autophagy in mouse hearts. J Appl Physiol (1985) 134, 1287–1299 (2023).

76 Guridi, M. et al. Activation of mTORC1 in skeletal muscle regulates whole-body metabolism through FGF21. Science signaling 8, ra113 (2015).

77 Zhang, I. X., Ren, J., Vadrevu, S., Raghavan, M. & Satin, L. S. ER stress increases store-operated Ca(2+) entry (SOCE) and augments basal insulin secretion in pancreatic beta cells. J Biol Chem 295, 5685–5700 (2020).

78 Castets, P., Ham, D. J. & Ruegg, M. A. The TOR Pathway at the Neuromuscular Junction: More Than a Metabolic Player? Front Mol Neurosci 13, 162 (2020).

79 Thornton, A. M. et al. Store-operated Ca(2+) entry (SOCE) contributes to normal skeletal muscle contractility in young but not in aged skeletal muscle. Aging (Albany NY) 3, 621–634 (2011).

80 Zhao, X. et al. Compromised store-operated Ca2+ entry in aged skeletal muscle. Aging Cell 7, 561–568 (2008).

81 Dubowitz, V. & Sewry, C. Muscle Biopsy: A Practical Approach. 4th edition edn, (Elsevier Health Sciences, 2013).

82 Moyle, L. A. & Zammit, P. S. Isolation, culture and immunostaining of skeletal muscle fibres to study myogenic progression in satellite cells. Methods Mol Biol 1210, 63–78 (2014).

