## Supplemental Data for "mTORC1-dependent SOCE activity regulates synaptic gene expression and muscle response to denervation"

Supplementary materials contain 7 Supplementary Figures, and 1 Supplementary Table.

**References**


1 Ham, D. J. *et al.* The neuromuscular junction is a focal point of mTORC1 signaling in sarcopenia. *Nat Commun* **11**, 4510 (2020).

2 Petrany, M. J. *et al.* Single-nucleus RNA-seq identifies transcriptional heterogeneity in multinucleated skeletal myofibers. *Nat Commun* **11**, 6374 (2020).

### Supplementary Figures

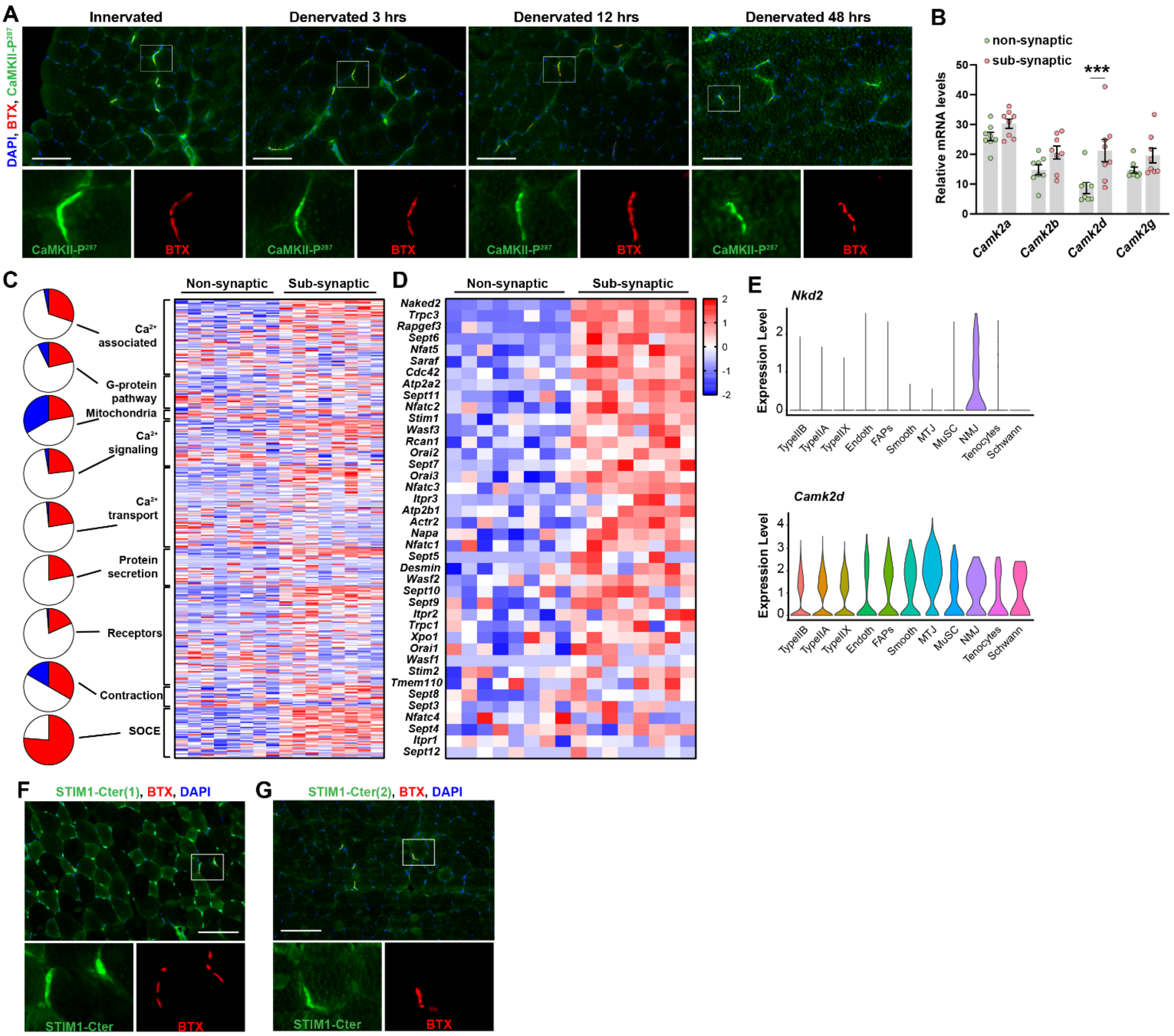


**Fig. S1. Ca$^{2+}$-related pathways at the endplate region.**

(**A**) Representative immunostaining for CaMKII-P$^{287}$ of innervated and denervated (3, 12 or 48 hrs) TA muscle sections. Scale bar, 100 µm. (**B**) mRNA levels of *Camk2* genes in non- and sub-synaptic regions (data from SarcoAtlas[1]). n = 8. Values are mean ± s.e.m; ***P < 0.001; unpaired two-tailed Student's t-test. (**C, D**) Heatmaps showing the expression of Ca$^{2+}$-related genes (**C**) or Store-Operated-Calcium-Entry (SOCE)-related genes (**D**) in non- and sub-synaptic regions of TA muscle (data from SarcoAtlas[1]). Each column represents an individual sample, each row represents a gene. Pie charts indicates the proportion of genes with a predominant expression in the sub-synaptic region (red), in the non-synaptic region (blue) or similarly express in both regions (white). (**E**) Violin plots showing expression levels of *Nkd2* and *Camk2d* in clusters of single nuclei isolated from TA muscle from 5-month-old mice [2]. Endoth: endothelial cells; FAP: Fibro/adipogenic progenitors; Smooth: smooth muscle; MTJ: myotendinous junction; MuSC: muscle stem cell. (**F, G**) Representative immunostaining of TA muscle sections for STIM1-Cter (with two different antibodies). The sub-synaptic region is visualized with α-Bungarotoxin (BTX). Scale bar, 100 µm.

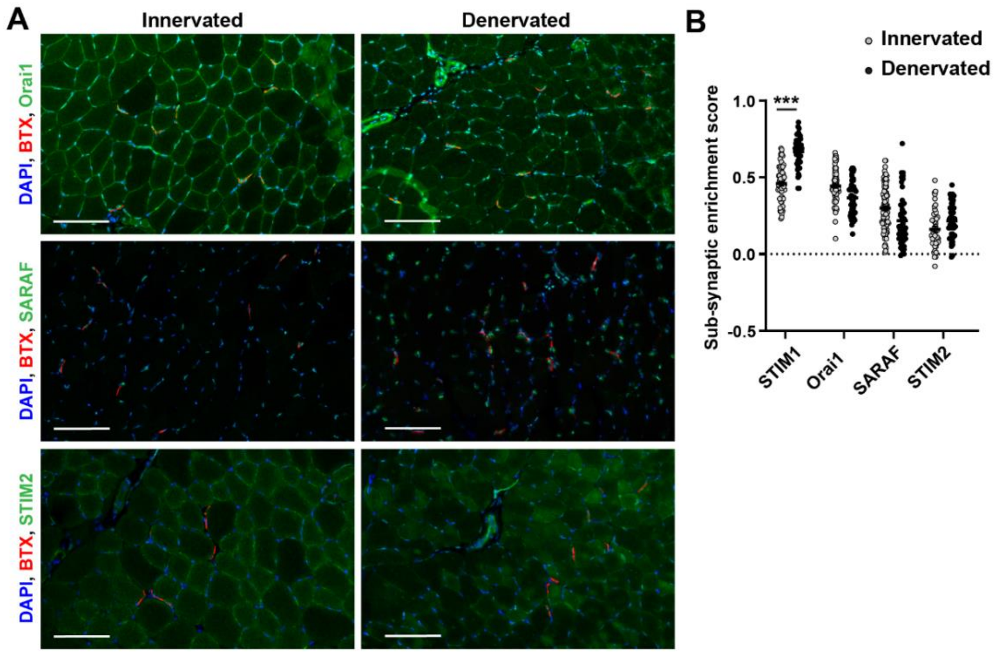


**Fig. S2. Localization of SOCE components in denervated muscle**

(**A**) Representative immunostaining of innervated and denervated (7 days) TA muscle sections for SOCE components, with BTX staining. Scale bar, 100 µm. (**B**) Sub-synaptic protein enrichment score based on the quantification of the colocalization of STIM1, Orai1, SARAF and STIM2 with BTX staining on muscle sections from innervated or denervated (7 days) TA muscles. Each dot represents an NMJ (> 40 NMJ from 4 independent samples). ***P < 0.001; unpaired two-tailed Student's t-test.

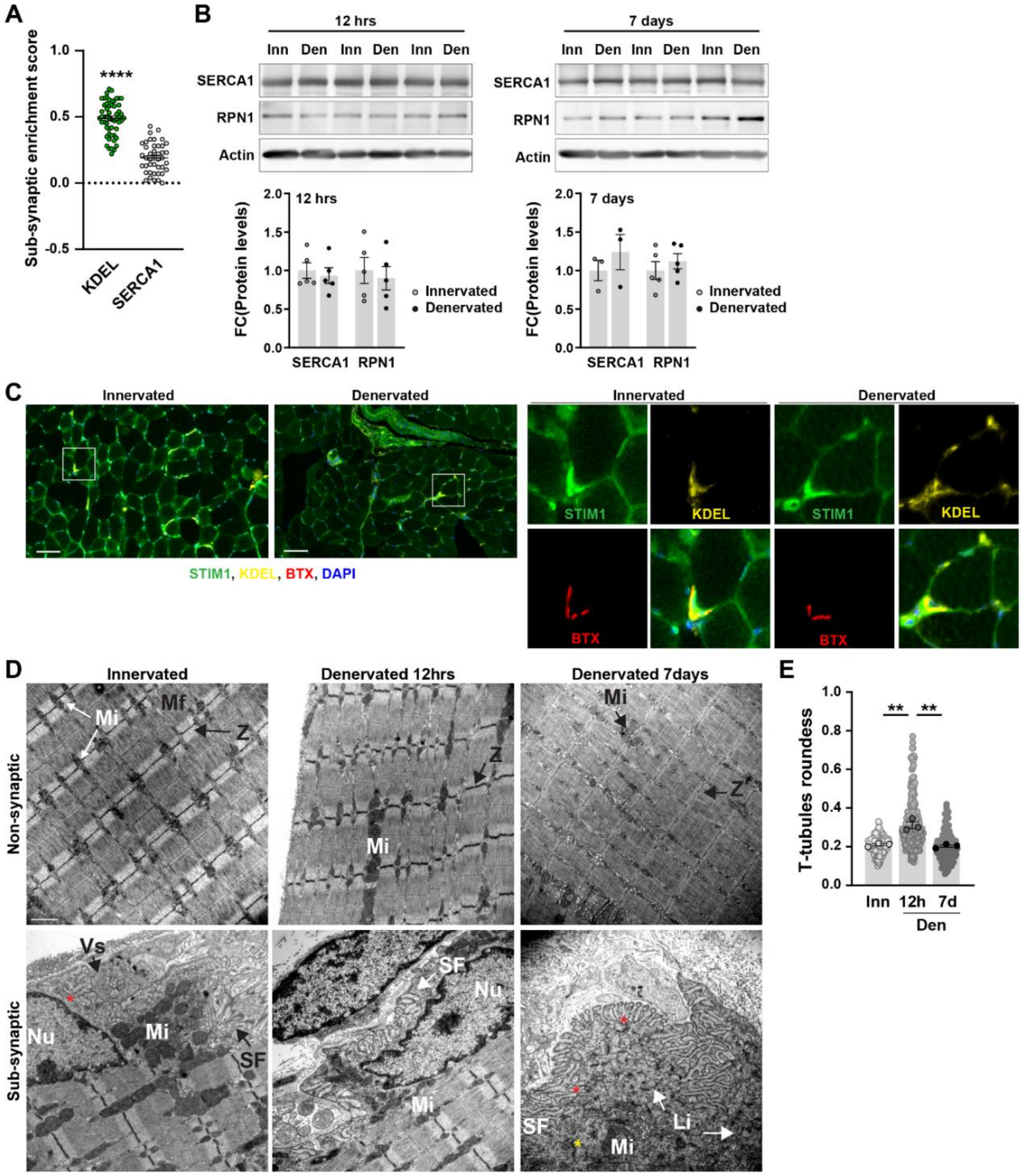


**Fig. S3. rER accumulates in the sub-synaptic muscle region.**

(**A**) Sub-synaptic protein enrichment score for SERCA1 and KDEL is based on the quantification of the colocalization of proteins and BTX staining, presented in Fig. 3, A and B. Each dot represents an NMJ (> 40 NMJ from 4 independent samples). (**B**) Representative immunoblots of SERCA1 and RPN1 in innervated (Inn) and denervated (Den; 12hrs or 7 days) TA muscles. Quantification of protein levels is relative to Actin levels. n=5 (except for SERCA1 7days, n=3). (**C**) Representative immunostaining for STIM1 and KDEL of innervated and denervated (7 days) TA muscle sections. BTX is used to visualize the sub-synaptic region. Scale bar, 100 µm. (**D**) Transmission electron microscopy of longitudinal sections of EDL innervated and denervated (Den, 12 hrs or 7 days) muscles, in non- and sub-synaptic regions. Scale bar, 1µm. Mi, Mf, Z, Vs, Nu, SF, Li indicate mitochondria, myofibrils, Z-lines, pre-synaptic vesicles, myonuclei, sarcoplasmic folds, and lipid droplets, respectively. Red and yellow asterisks point to rER contact points with plasma membrane or mitochondria, respectively. (**E**) T-tubules roundness quantified in innervated (Inn) and denervated (Den, 12 hrs or 7 days) EDL muscles based on electron micrographs. Light and opaque dots represent a T-tubule and the mean for one mouse, respectively. n=3. All values are mean ± s.e.m; **$P < 0.01$, ****$P<0.0001$; P value calculated from t score (A, see methods for more details), one-way ANOVA with Benjamini and Hochberg's post-hoc analysis to correct for multiple comparisons (E).

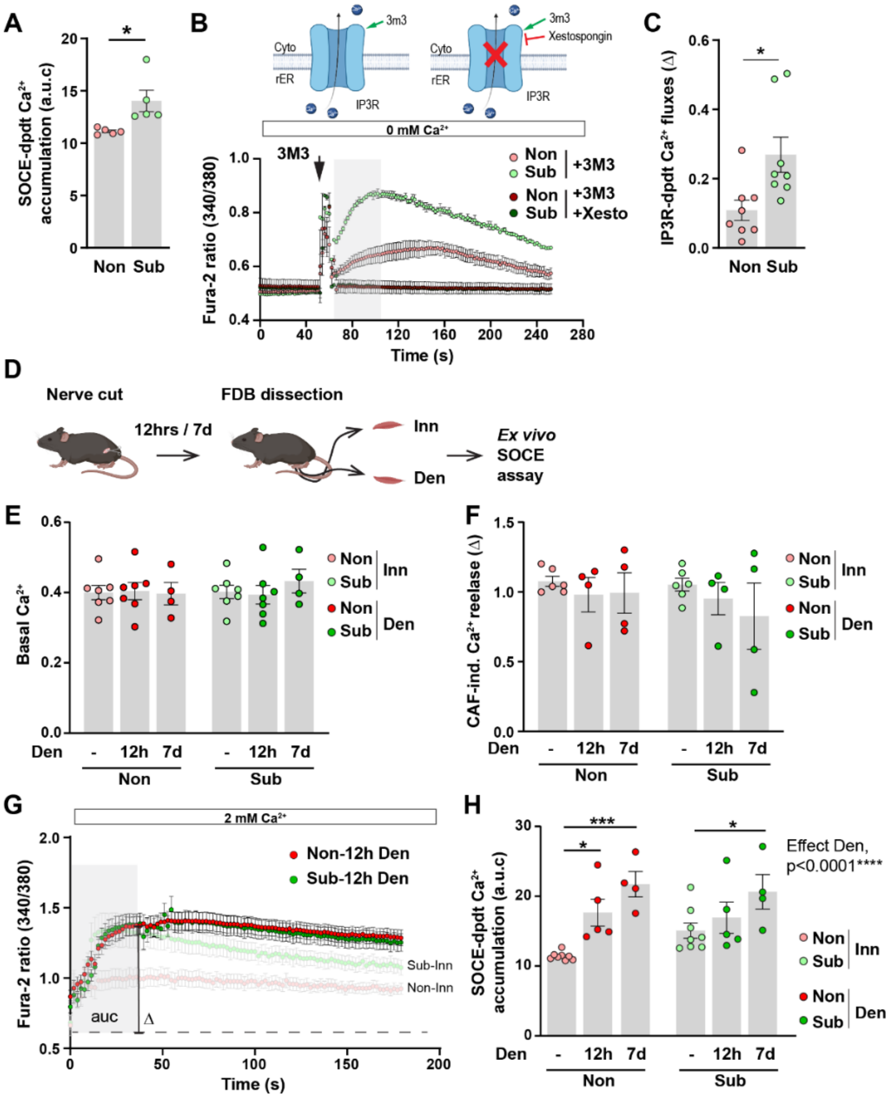


**Fig. S4. Denervation-induced changes in $Ca^{2+}$ fluxes**

(**A**) SOCE-dependent $Ca^{2+}$ entry measured as the area under the curve to the peak (a.u.c) in non- and sub-synaptic regions of innervated muscle fibers. n=5. (**B**) Representative cytosolic $Ca^{2+}$ traces in non- and sub-synaptic regions of isolated FDB fibers in response to m-3M3-FBS, with (+Xesto) or without Xestospongin C, in the absence of $Ca^{2+}$ in the medium. (**C**) Amplitude of IP3R-induced $Ca^{2+}$ fluxes in non- and sub-synaptic regions of isolated FDB fibers. n=8. (**D**) Workflow for the analysis of SOCE in innervated and denervated FDB muscle fibers. (**E, F**) Quantification of cytosolic basal $Ca^{2+}$ concentration (**E**) and caffeine (CAF)-induced $Ca^{2+}$ release (**F**) in non- and sub-synaptic regions of innervated and denervated (Den, 12hrs or 7 days) muscle fibers. (**G, H**) Mean SOCE traces (**G,** Den, 12 hours) and SOCE-dependent $Ca^{2+}$ accumulation (**H,** Den, 12 hours and 7 days) in non- and sub-synaptic regions of innervated and denervated FDB muscle fibers. In **G**, light traces correspond to innervated fibers (Inn). n=8 (Inn), 5 (Den, 12 hours), 4 (Den, 7 days). All values are mean ± s.e.m; *P < 0.05, ***P < 0.001; unpaired two-tailed Student's t-test (A, C), two-way ANOVA with Benjamini and Hochberg's post-hoc analysis to correct for multiple comparisons (H).

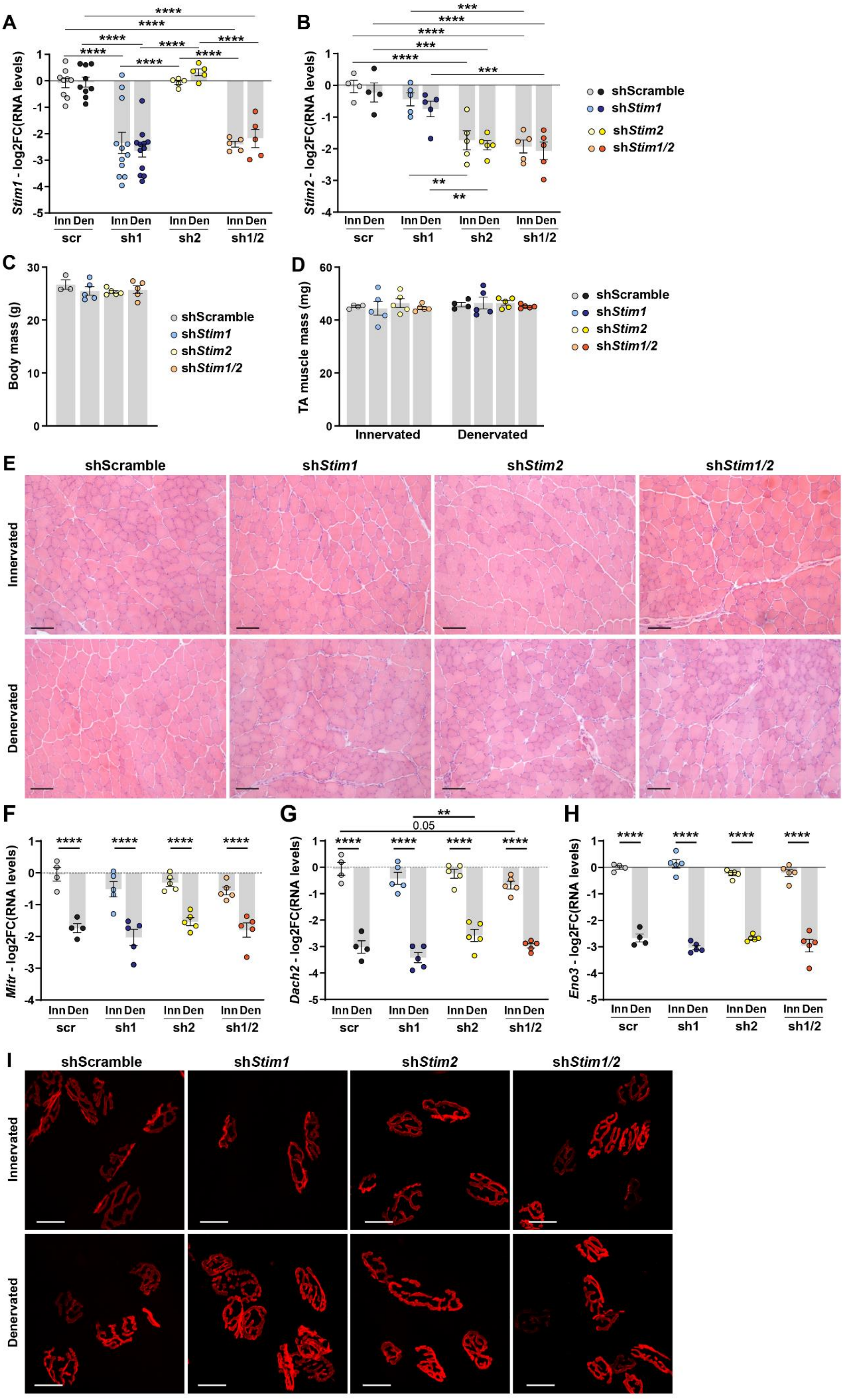
A
B
C
D
E
F
G
H
I
shScramble
shStim1
shStim2
shStim1/2
Stim1 - log2FC(RNA levels)
Stim2 - log2FC(RNA levels)
Inn Den
scr
sh1
sh2
sh1/2
Body mass (g)
TA muscle mass (mg)
Innervated
Denervated
Mitr - log2FC(RNA levels)
Dach2 - log2FC(RNA levels)
Eno3 - log2FC(RNA levels)
0.05

**Fig.S5. SOCE perturbation alters muscle response to denervation.**

(**A, B**) mRNA levels of *Stim1* (**A**) and *Stim2* (**B**) in innervated or denervated (3 days) TA muscles infected with AAV-shScramble, -sh*Stim1*, -sh*Stim2*, or –sh*Stim1/Stim2*. Levels are normalized on *Tbp* and expressed as log2(fold change) of control innervated muscle injected with AAV-shScramble. n=9scr, 12sh1, 5sh2, 5sh1/2 (A); 4scr, 5sh1, 5sh2, 5sh1/2 (B). (**C**) Body mass of mice injected with AAV-sh-Scramble, -sh*Stim1*, -sh*Stim2*, or –sh*Stim1/Stim2*. n=5 (except for Scr, n=3). (**D**) Mass of innervated and denervated TA muscles infected with AAV-shScramble, -sh*Stim1*, -sh*Stim2*, or –sh*Stim1/Stim2*. n=5 (except for scr, n=4). (**E**) Hematoxylin and eosin staining of innervated and denervated TA muscles injected with AAV-shScramble, -sh*Stim1*, -sh*Stim2*, or –sh*Stim1/Stim2*. Scale bar, 200 µm. (**F-H**) mRNA levels of *Mitr* (**F**), *Dach2* (**G**) and *Eno3* (**H**) in innervated and denervated (3 days) TA muscles infected with AAV-sh-Scramble, -sh*Stim1*, -sh*Stim2*, or –sh*Stim1/Stim2*. RNA levels are normalized to *Tbp* and expressed as log2(fold change) of control innervated muscle injected with AAV-shScramble. n=4scr, 5sh1, 5sh2, 5sh1/2. (**I**) Representative images of muscle bundles stained with BTX isolated from EDL muscles infected with AAV-shScramble, -sh*Stim1*, -sh*Stim2*, or –sh*Stim1/Stim2*. Scale bars, 20 µm. All values are mean ± s.e.m; **P < 0.01, ***P < 0.001, ****P < 0.0001; two-way ANOVA with Benjamini and Hochberg's post-hoc analysis to correct for multiple comparisons (A, B, F-H).

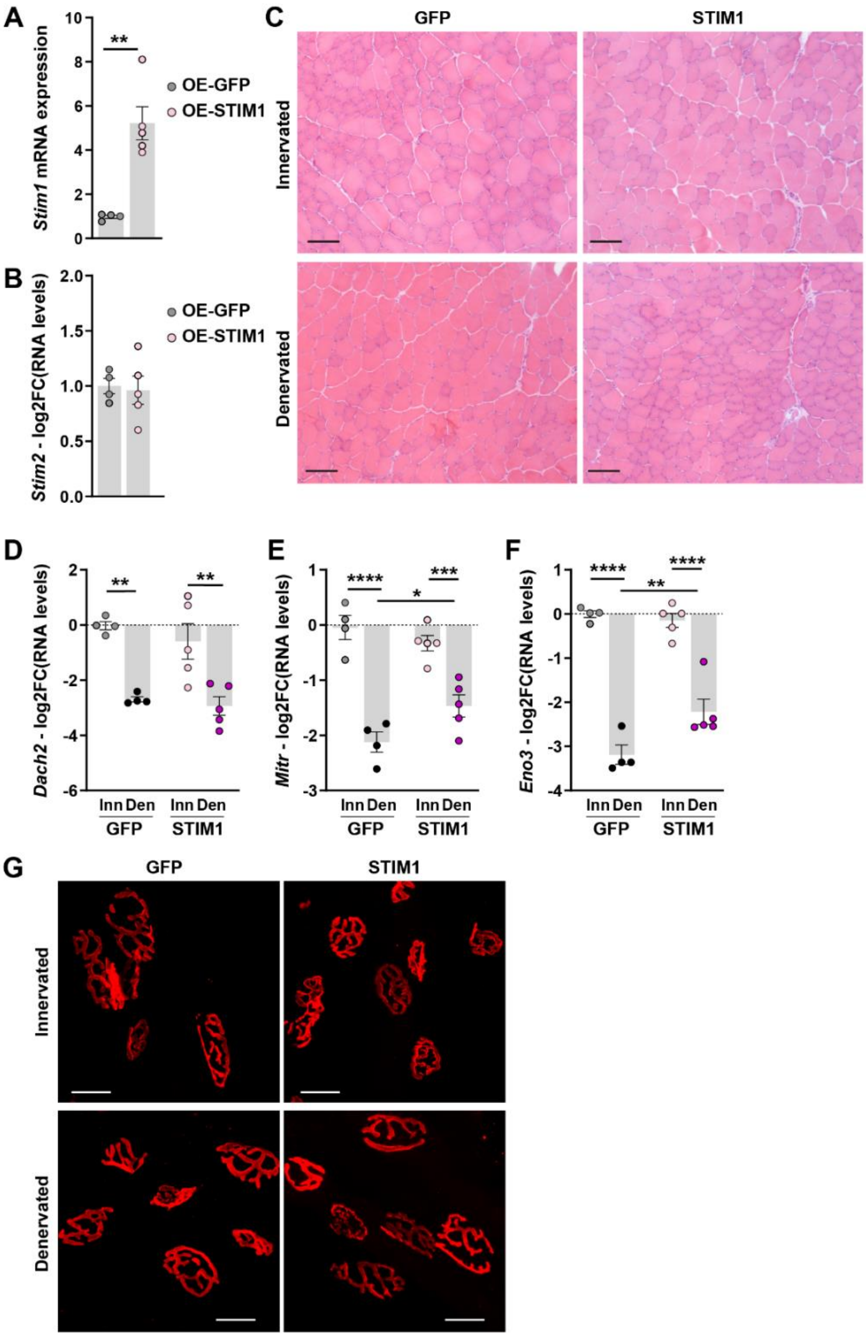


**Fig. S6. STIM1 overexpression impairs muscle sensing of innervation/denervation.**

(**A, B**) mRNA levels of *Stim1* (**A**) and Stim2 (**B**) in innervated or denervated (3 days) TA muscles infected with AAV-eGFP or -STIM1. Levels are normalized on *Tbp* and expressed as log2(fold change) of control innervated infected with AAV-eGFP. n=4GFP/5STIM1. (**C**) Hematoxylin and eosin staining of innervated and denervated TA muscles injected with AAV-eGFP or –STIM1. Scale bar, 200 µm. (**D-F)** mRNA levels of *Dach2* (**D**), *Mitr* (**E**) and *Eno3* (**F**) in innervated and denervated (3 days) TA muscles infected with AAV-eGFP or -STIM1. Levels are normalized on *Tbp* and expressed as log2(fold change) of control innervated muscle injected with AAV-eGFP. n=4Inn/5Den. (**G**) Representative images of muscle bundles stained with BTX isolated from EDL muscles infected with AAV-eGFP or -STIM1. Scale bar, 20 µm. All values are mean ± s.e.m; * $P<0.05$, ** $P < 0.01$, *** $P < 0.001$, **** $P < 0.0001$; unpaired two-tailed Student's t-test (A, B), two-way ANOVA with Benjamini and Hochberg's post-hoc analysis to correct for multiple comparisons (D-F).

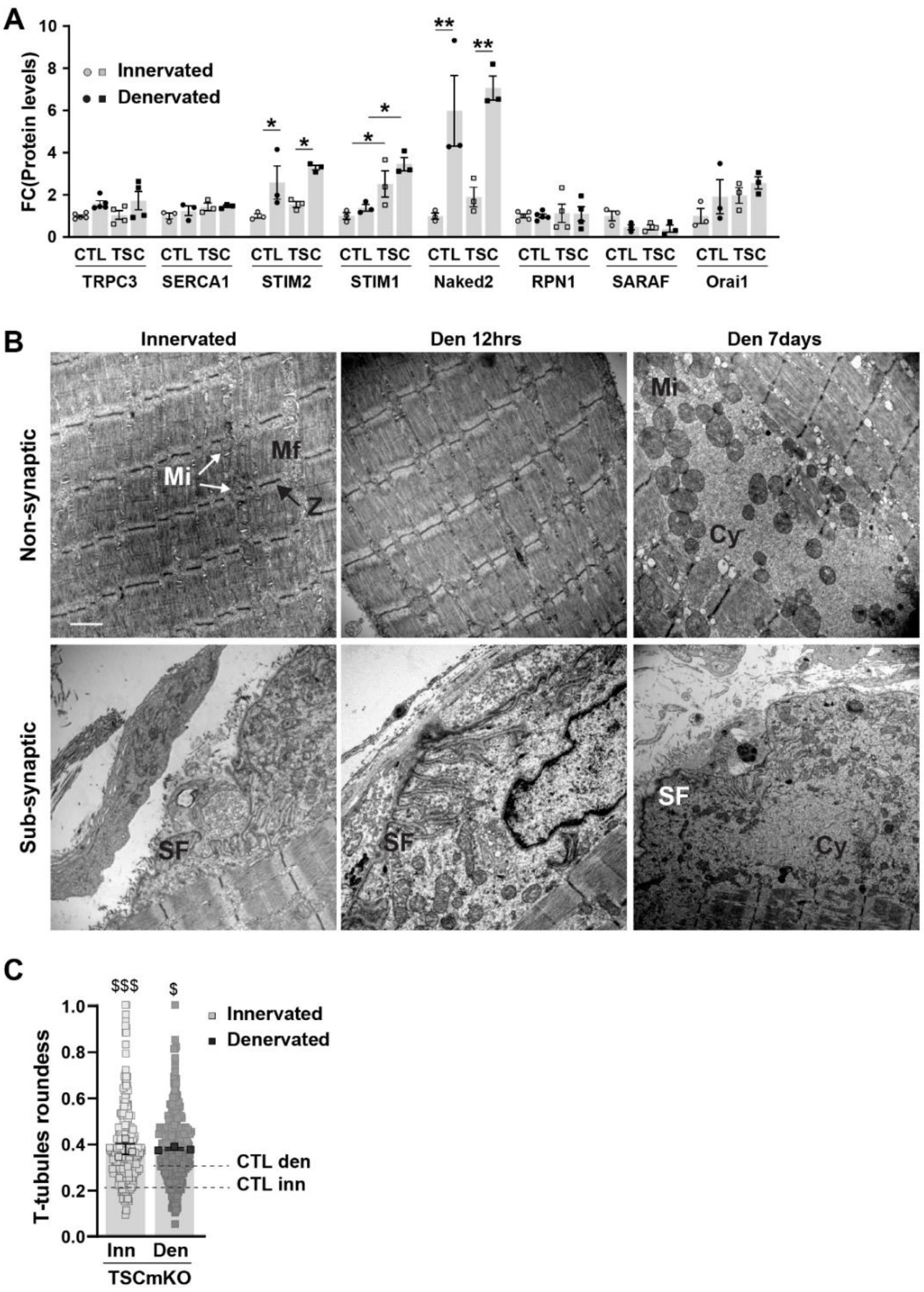


**Fig. S7. mTORC1 activation perturbs SOCE pathway and muscle response to denervation.**

(**A**) Levels of proteins involved in SOCE in innervated (Inn) and denervated (Den, 7 days) TA muscles from control (CTL) and TSCmKO (TSC) mice. Levels are relative to Actin and normalized on control innervated muscle. n=3 (except for TRPC3 and RPN1, n=5CTL/4TSC). (**B**) Transmission electron microscopy of longitudinal sections of TSCmKO EDL innervated and denervated (Den, 12 hrs or 7 days) muscles in non- and sub-synaptic regions. Scale bar, 1µm. Mi, Mf, Z, Cy, SF indicate mitochondria, myofibrils, Z-lines, cytoplasmic space, and sarcoplasmic folds, respectively. (**C**) T-tubules roundness quantified in innervated (Inn) and denervated (Den, 12 hrs) EDL muscles from

TSCmKO mice, based on electron micrographs. Light and opaque dots represent a T-tubule and the mean for one mouse, respectively. Dashed lines indicate mean values of control mice (CTL). n=3. All values are mean ± s.e.m; *P < 0.05, **P<0.01, between Inn/Den; $^{\$}$P < 0.05, $^{\$\$\$}$P < 0.001, between genotypes; two-way ANOVA with Benjamini and Hochberg’s post-hoc analysis to correct for multiple comparisons (A, C).

**Supplementary Table S1 | Sequence of qPCR primers used in this study.**

| Gene | Forward primer | Reverse primer |
|---|---|---|
| *Chrna1* | TCCCTTCGATGAGCAGAACT | GGGCAGCAGGAGTAGAACAC |
| *Chrng* | GTGTCTTCGAGGTGGCTCTC | ACAGAGATGGAGCAGGAGGA |
| *Dach2* | CCAGCTCAAATCCCAGTCAT | CGCAGTTCCTTCTTTTCCTG |
| *Hdac4* | CAGACAGCAAGCCCTCCTAC | AGACCTGTGGTGAACCTTGG |
| *Mitr* | CCACCTTGAAGAAGCAGAGG | TGGTGTCCTTAGAGGCTGCT |
| *Mse* | GGGAGATGACCTCACGGTAA | TTACAGGCCTGGATGGACTC |
| *Myog* | CACTCCCTTACGTCCATCGT | CAGGGCTGTTTTCTGGACAT |
| *Stim1* | ACGATGCCAATGGTGATGTG | CACCTCATCCACAGTCCAGT |
| *Stim2* | AGATGGACGACGACAAGGAC | CAGGGTATCCTCAAGTGTCCA |
| *Tbp* | CTCAGTTACAGGTGGCAGCA | CAGCACAGAGCAAGCAACTC |
